# Does insect herbivory suppress ecosystem productivity? Evidence from a temperate woodland

**DOI:** 10.1101/2021.01.28.428605

**Authors:** Kristiina Visakorpi, Sofia Gripenberg, Yadvinder Malhi, Terhi Riutta

**Affiliations:** Department of Zoology, University of Oxford, Oxford, UK; School of Biological Sciences, University of Reading, Reading, UK; Environmental Change Institute, School of Geography and the Environment, University of Oxford, Oxford, UK; Imperial College London, Department of Life Sciences, Silwood Park Campus, Ascot, UK

**Keywords:** insect herbivory, carbon cycle, net primary productivity, gross primary productivity, respiration, *Quercus robur*, *Operophtera brumata*, Wytham Woods

## Abstract

Our current understanding of the relationship between insect herbivory and ecosystem productivity is limited. Previous studies have typically quantified only leaf area loss, or have been conducted during outbreak years. These set-ups often ignore the physiological changes taking place in the remaining plant tissue after insect attack, or may not represent typical, non-outbreak herbivore densities. Here, we estimate the amount of carbon lost to insect herbivory in a temperate deciduous woodland both through leaf area loss and, notably, through changes in leaf gas exchange in non-consumed leaves under non-outbreak densities of insects. We calculate how net primary productivity changes with decreasing and increasing levels of herbivory, and estimate what proportion of the carbon involved in the leaf area loss is transferred further in the food web. We estimate that the net primary productivity of an oak stand under ambient levels of herbivory is 54 - 69% lower than that of a completely intact stand. The effect of herbivory quantified only as leaf area loss (0.1 Mg C ha^−1^ yr^−1^) is considerably smaller than when the effects of herbivory on leaf physiology are included (8.5 Mg C ha^−1^ yr^−1^). We propose that the effect of herbivory on primary productivity is non-linear and mainly determined by changes in leaf gas exchange. We call for replicated studies in other systems to validate the relationship between insect herbivory and ecosystem productivity described here.

## Introduction

By affecting plant abundance, distribution and physiology, herbivores can have large impacts on ecosystem carbon storage and cycling (Chapin 1997, Kurz et al. 2008, Clark et al. 2010, Schäfer et al. 2010, Heliasz et al. 2011, Estes et al. 2011, Schmitz et al. 2014, Flower and Gonzalez-Meler 2015, Lund et al. 2017). To date, many studies have demonstrated the role of large grazers and their predators in controlling primary productivity (e.g. Zimov et al. 2009, Estes et al. 2011, Schmitz et al. 2014, Wilmers and Schmitz 2016) or insect outbreaks in causing defoliation and tree mortality (Kurz et al. 2008, Clark et al. 2010, Amiro et al. 2010, Edburg et al. 2012, Flower et al. 2013, Flower and Gonzalez-Meler 2015). Under outbreak densities, the effects of insect herbivores on forest carbon cycling can be large enough to shift the ecosystem from being a carbon sink into a carbon source (Kurz et al. 2008, Heliasz et al. 2011, Lund et al. 2017).

However, the role of non-outbreak, low-density populations of insect herbivores on carbon cycling has received less attention. The ubiquity of insect herbivory in terrestrial ecosystems (Strong et al. 1984, Schoonhoven et al. 2005, Forister et al. 2015), and its potential effects on the rates of photosynthesis and respiration of the remaining plant tissue (Nykänen and Koricheva 2004, Nabity et al. 2009, Bilgin et al. 2010) suggest that even low densities of insect herbivores could have a large impact on ecosystem carbon sequestration and loss (Strickland et al. 2013, Visakorpi et al. 2018).

Although insect herbivory has been shown to affect rates of photosynthesis and plant respiration beyond leaf area loss (“indirect effect of herbivory”, as opposed to “direct effect” of leaf area loss; Oleksyn et al. 1998, Zangerl et al. 2002, Nykänen and Koricheva 2004, Nabity et al. 2009, Bilgin et al. 2010, Meza-Canales et al. 2017), these physiological plant responses to herbivory have generally been neglected when estimating changes in ecosystem CO_2_ exchange. Consequently, we currently do not have a clear picture how the most typical levels of insect herbivory affect ecosystem CO_2_ exchange. Ignoring the effects of insect herbivory could lead to biased estimations of ecosystem productivity, or of the role of forests as carbon sinks (Kurz et al. 2008, Campioli et al. 2016).

In this study, we quantify the effects of insect herbivory on forest-level net primary productivity in a temperate maritime woodland in southern England, UK. In a previous study in this system (Visakorpi et al. 2018), we found that photosynthesis of oak (*Quercus robur*, L.) was substantially lower in leaves subjected to herbivory by winter moth (*Operophtera brumata*, L.) caterpillars than in intact leaves surrounded only by other intact leaves. Moreover, a similar reduction in photosynthetic rate was seen in intact leaves on the same shoots as the damaged leaves, resulting in an estimated 50% reduction in canopy-level photosynthesis. How these changes in carbon assimilation affect tree and stand level productivity remains unknown. Here, we combine our previous measurements of herbivory-inflicted leaf area loss and indirect changes in photosynthetic rate (Visakorpi et al. 2018) with measurements on canopy and woody respiration, hourly meteorological data and tree survey data from the study site to estimate the effects of herbivory on oak primary production over a growing season (Figure 1). We ask: 1) what is the potential effect of insect herbivory on net primary production (NPP) of an oak stand, estimated separately through leaf area loss and through changes in canopy gas exchange and 2) how does the effect of herbivory on forest productivity change with changing herbivory pressure?

**Figure 1.**
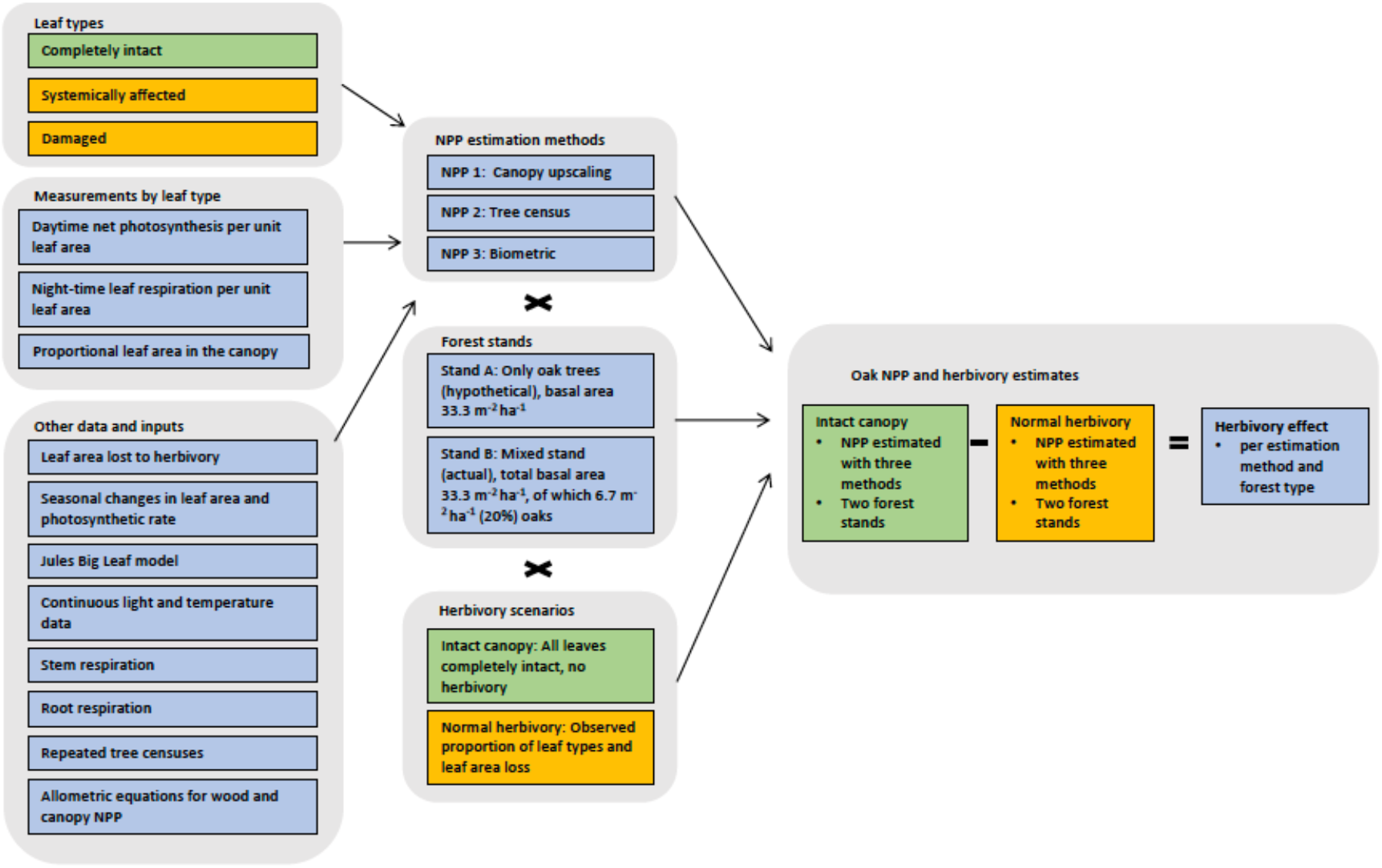
The summary of the data inputs and methods used to estimate NPP and the effect of herbivory.

## Methods

### Study site and system

All data were collected in Wytham Woods (51°.47′ N, 1° 20′ W, 160 m.a.s.l), located in Oxfordshire, UK. Wytham Woods is an ancient semi-natural woodland of 390 ha. Most parts of the woodland have experienced long-standing management practices (e.g. coppicing, selective logging) until 50 year ago (Savill 2011). Recruitment is low, and established vegetation accounts for most of the carbon uptake (Thomas et al. 2011). The area within Wytham Woods in which our study took place has been a woodland at least since the 18^th^ century, and most likely since the last ice age (Savill 2011).

Our focal study area is an 18-ha forest dynamics monitoring plot, part of the Smithsonian Global Earth Observatory network (www.forestgeo.si.edu). The plot is dominated by sycamore (*Acer pseudoplatanus*, ca. 50% of the basal area), pedunculate oak (*Quercus robur*, ca. 20% of the basal area) and ash (*Fraxinus excelsior*, ca. 20% of the basal area; Butt et al. 2009, Savill 2011). The plot was surveyed for tree growth, recruitment and mortality in 2008, 2010 and 2016. The carbon cycle of the plot has been measured using the eddy covariance technique (Thomas et al. 2011) and biometric quantification of leaf and woody biomass in a 1 ha subplot (Fenn et al. 2015, Table 1). Previous work at the site has revealed seasonal dynamics in oak photosynthesis, which peaks on average 63 (± 6) days after budburst (Morecroft et al. 2003).

**Table 1.**
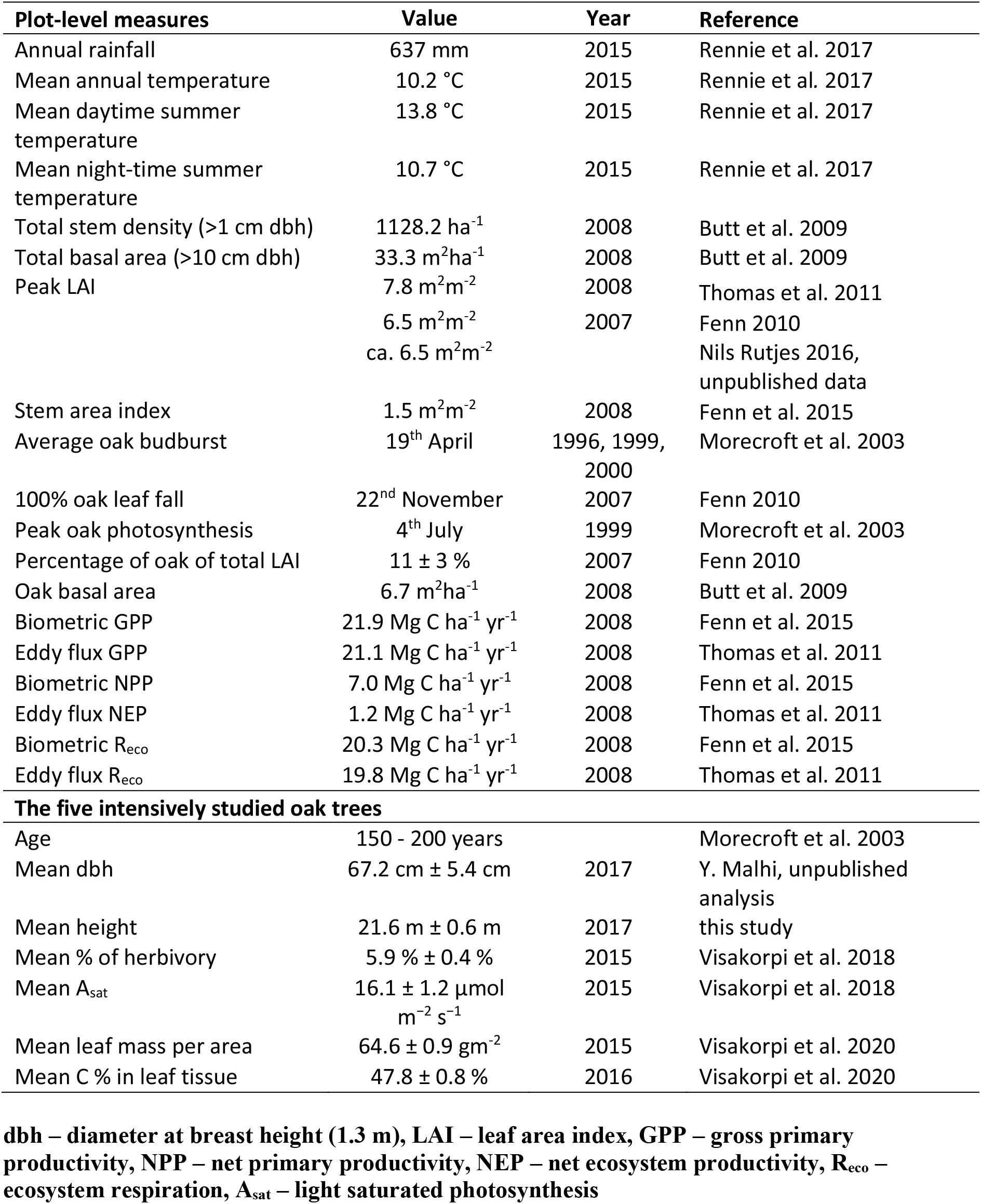
Characteristics of the study site, and the five oak trees used for the gas exchange measurements and herbivory manipulations. Errors are ± 1 SEM.

In Wytham Woods, lepidopterous caterpillars are the first herbivores to attack the newly flushed leaves of most tree species (Feeny 1970). They emerge in synchrony with the budburst, and feed until early June (Hunter 1992). Relatively few herbivore species feed on the mature leaves later in the season (Feeny 1970). Of the early-spring herbivores, the winter moth (*Operophtera brumata*) is one of the most common species (Feeny 1970), feeding on most of the trees in the area. The winter moth is native to the area, and its population reaches outbreak densities approximately every 10 years. The last high abundance year prior to our study was in 2010 (285 caterpillars per m^2^ of oak canopy, Lionel Cole; unpublished data).

Since 2013, the population size has been small (e.g. 5 caterpillars per m^2^ in 2014; Lionel Cole, unpublished data).

### Quantifying leaf area loss and the effect of insect herbivory on leaf-level photosynthesis

Data on leaf area loss were collected from leaves in the upper canopy of five mature oak trees located in the 18-ha plot, accessible from a canopy platform. To estimate the level of insect herbivory on the five focal oaks throughout the growing season, we used estimates of the proportion of leaf area loss and the frequency distributions of three different leaf types per tree: completely intact leaves on shoots with only intact leaves (“completely intact”), intact leaves on shoots with at least one herbivore-damaged leaf (“systemically affected”), and leaves damaged by herbivores (“damaged”) (Visakorpi et al. 2018). Data on the effect of winter moth caterpillars on leaf-level photosynthesis were obtained from the manipulative experiment reported in Visakorpi et al. (2018). In brief, we introduced winter moth caterpillars on selected oak shoots, and subsequently measured light-photosynthesis response curves on three leaf types (intact, systemically affected, damaged). To estimate the amount of carbon lost through the consumed leaf tissue, we used estimates for leaf mass per area (“LMA”, gm^−2^) and leaf carbon content (% of dry mass) for the same leaf types from the same experiment (Visakorpi et al. 2020; Appendix 2, Table S3).

#### Estimating oak NPP

Figure 1 summarizes our approach and Appendix 1 provides more detail on the methodology. In brief, we used three methods to estimate oak NPP. First, NPP was estimated as the difference between canopy net photosynthesis and woody (stem + root) respiration (“*NPP through canopy upscaling*”). Second, we used census data on woody growth and combined these data with allometric equations to estimate leaf production and belowground production (“*NPP through tree growth census*”). Third, we used earlier estimates of oak above- and belowground NPP at the site obtained from biometric measurements (“*NPP through biometric measurements*”) (Fenn 2010, Fenn et al. 2015). For each of the three methods, we estimate oak NPP for a hypothetical stand comprising only of mature oak trees, assigning the total basal area (33 m^2^ ha^−1^) of trees at the site to oaks only, and for the actual per ha of oak of the site (oak basal area 6.7 m^2^ ha^−1^). For each case, we estimate oak NPP for two scenarios: 1) for an intact canopy (hereafter “*Intact canopy”*) and 2) for a canopy under the observed level of herbivory (*“Normal herbivory”*). The effect of insect herbivory on NPP is the difference in NPP between the *Intact canopy* and the *Normal canopy* scenarios.

#### NPP through canopy upscaling

We used photosynthesis-light response curves measured from completely intact, systemically affected and damaged leaves from the five focal oak trees (one per leaf per leaf type per tree, Table 1). The leaf-level measures were scaled up with the Big Leaf approach of The Joint UK Land Environment Simulator (“JULES”, Clark et al. 2011). The canopy assimilation model included estimates for photosynthetic rate and daytime respiration rate (i.e. net photosynthesis) scaled to changes in LAI through the canopy, while taking into account leaf area loss to herbivory and the frequency distributions of the three different leaf types. Light was assumed to be reduced through the canopy layers (Monsi and Saeki 1953), photosynthesis to respond to light according to the light-photosynthesis response curves, and increased light to inhibit respiration (Mercado et al. 2007). To estimate night-time respiration, we measured respiration rates on the three leaf types on each of our five focal trees during two nights in July 2015. To scale night respiration to canopy level, we accounted for the frequency distributions of different leaf types and corrected for differences in respiration rates between different canopy layers (Griffin et al. 2001). Canopy gas exchange (daytime net photosynthesis – night respiration) was scaled to hourly light (daytime net photosynthesis) and temperature (night respiration) data, to seasonal changes in LAI (Stokes 2000, Fenn 2010) and to seasonal changes in photosynthetic rate of oak (Morecroft et al. 2003). The canopy night respiration rate was further scaled with the daytime photosynthetic rate of the previous day (Whitehead et al. 2004). To estimate woody respiration, we used data from previous measurements of oak stem respiration at the site (Walker 2017). We assumed that stem respiration followed the same pattern as leaf respiration in response to herbivory, since both leaf and stem respiration rates have a similar, positive relationship with photosynthesis (Wertin and Teskey 2008). Stem respiration was scaled to hourly temperature data. Root respiration was assumed to be 13% of stem respiration, based on earlier measurements at the site (Fenn et al. 2015). Canopy gas exchange and woody respiration were scaled to plot level by assuming leaf area index (“LAI”) of 6.5 m^2^/m^2^ and stem area index (“SAI”) of 1.5 m^2^/m^2^, both previously estimated for the site (Fenn 2010, Nils Rutjes 2016, unpublished data; Fenn and others 2015, Table 1). For all meteorological data, we used hourly data recorded at an open site in Wytham Woods during years 2014 and 2015 (Figure 2, Rennie et al. 2017).

**Figure 2.**
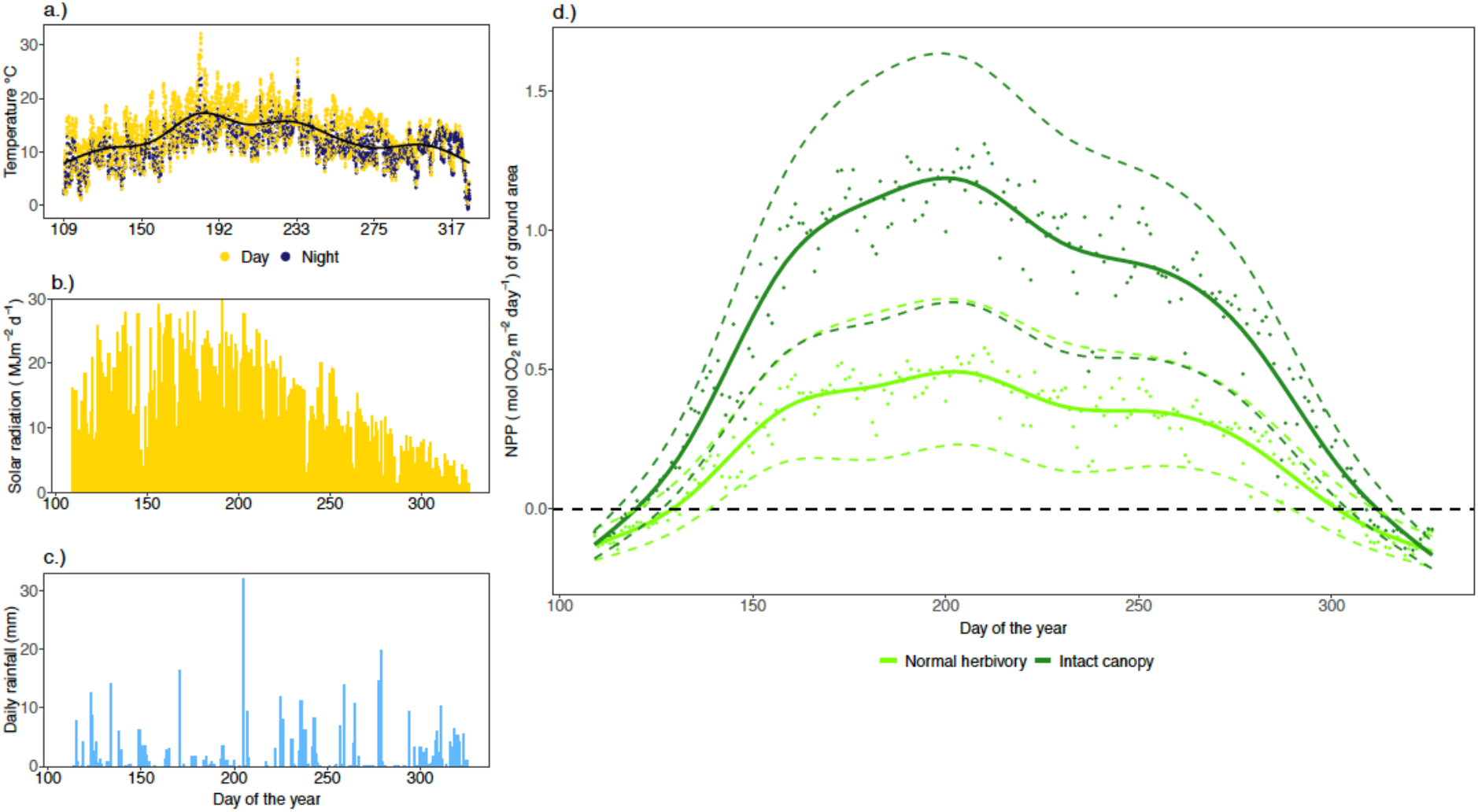
Meteorological data and net primary productivity in Wytham Woods. a) Hourly temperature for the growing season 2015 (19th April – 22nd November), which were used for upscaling stem, root and night-time canopy respiration. The average temperature was 13.8 °C during the day and 10.7 °C during the night. b) Solar radiation per day over the season 2014, which was used to upscale the canopy photosynthesis measurements. c) Amount of rainfall per day over the growing season 2015 (not used in the data analyses or upscaling). The annual rainfall (637 mm) was slightly less than the 1993–2010 average of 714 mm (Fenn et al. 2015). d) Net primary productivity (NPP) per m^2^ of ground area based on data from five oak trees during the growing season 2015. The “intact canopy” scenario is based on measurements on intact leaves surrounded by only intact leaves, whereas the estimates under “normal herbivory” are based on photosynthesis and respiration rates of three different leaf types (intact, damaged, systemically affected), weighted with their observed abundance. Each data point is a modelled estimate for the average rate of the five trees for one day. The dashed black line shows the level of zero NPP (negative values indicate that the tree is losing carbon). The solid coloured line shows a general additive model across the five trees, and the dashed coloured lines show the propagated measuring uncertainty.

#### NPP through tree growth census

We used dbh measurements from all oak trees (355 stems, >10 cm dbh) within the 18 ha forest dynamics plot obtained from censuses in 2010 and 2016 (Y. Malhi, unpublished analysis). We estimated the yearly growth as an increase in aboveground woody biomass divided by the length of the census interval. To estimate woody NPP we multiplied the estimated change in woody biomass by the carbon content of oak wood (0.47; Butt et al. 2009). We then used oak-specific NPP allocation patterns obtained from (Fenn 2010) to estimate oak leaf NPP and belowground NPP (Appendix 1, Table S2). To estimate the effect of herbivory on oak NPP, we estimated NPP in the absence of herbivory assuming that the total NPP under normal herbivory situation was reduced to the same extent (ca. 56%, Appendix 1, Table S2) as canopy gas exchange in the canopy upscaling calculations described above. We then extrapolated the values per tree to estimates per hectare of a plot with only oak trees, and per hectare of the actual study site.

#### NPP through biometric measurements

We combined data on aboveground oak NPP collected in a one hectare subplot of the 18 ha forest dynamics plot with information on the oak-specific ratios of wood NPP to leaf NPP and aboveground biomass to belowground biomass (Fenn 2010; Fenn et al. 2015; see Appendix 1, Table S2). The biometric estimates were based on dendrometer measurements (woody production), measures of leaf production (litter traps) and of root production (soil respiration and inputs). We combined these relationships for an estimate of whole oak NPP as Mg C yr^−1^ per ha of the actual study site. To estimate the effect of herbivory on oak NPP, we again assumed that the total NPP was reduced to the same extent as canopy gas exchange in the canopy upscaling calculations above.

### Predicting the effects of herbivory on canopy assimilation under varying levels of herbivory

To predict the effect of herbivory on oak assimilation under different levels of caterpillar herbivory, we first simulated changes in the proportions of the three different leaf types (intact, systemically affected, damaged) in the oak canopy. We assumed that the ratio of damaged leaves to systemically affected leaves (1:2.3) and leaf area loss per damaged leaf (8.53%) stay at the same levels as observed in our herbivory surveys until all shoots have at least one damaged leaf. After this, the proportion of damaged leaves was set to increase until all leaves were damaged. Finally, we set the proportion of leaf area loss per leaf to increase, until the tree was completely defoliated. In other words, we assumed that the herbivory first spreads evenly to all shoots, then to all leaves, and then increases per leaf. LAI was assumed to decrease as leaf area loss increased, increasing the amount of light reaching lower canopy levels (Figure S7, Appendix 2). We assumed a linear relationship between winter moth caterpillar density and leaf area loss, with 5 individuals m^−2^ in 2015 (Lionel Cole, unpublished data) corresponding to 5.9% leaf area loss (across the canopy) and to peak-season LAI of 6.5, and maximum reported density of 1200 individuals m^−2^ corresponding to complete defoliation (Feeny 1970). We then estimated daytime canopy net photosynthesis with herbivory levels ranging from 0% to 100% leaf area loss. To test how sensitive the estimated relationship between the level of herbivory and its effect on canopy assimilation was to the assumptions of our models, we estimated the same relationship under four alternative scenarios: 1) the difference in photosynthetic rate between intact and damaged leaves is smaller than our field measurements suggest, 2) the photosynthetic rate of intact leaves increases with increasing level of herbivory in the canopy, 3) herbivory spreads through the canopy following a different spatial pattern than in our initial assumptions and 4) a second leaf flush compensates for some of the leaf area loss. For details, see Appendix 3.

## Results and discussion

We estimate that insect herbivores remove on average 0.12 (± 0.02) Mg C ha yr^−1^ in an oak stand through leaf area loss. Through reducing photosynthesizing leaf area and the photosynthetic rate of intact leaf tissue, insect herbivores prevent between 8.5 (± 5.1, canopy upscaling) and 4.1 (± 2.1, tree census) Mg C ha^−1^ yr^−1^ of carbon from being assimilated (Figure 2 and 3; Appendix 2, Figure S5). Depending on the method, we estimate the NPP of a forest consisting of only mature oak trees to be between 3.4 (± 0.6; biometric measurements) and 4.6 (± 1.1; tree census) Mg C ha^−1^ yr^−1^ under the observed level of herbivory (5.9% leaf area loss, Table 2, Figure 3). This is between 54 ± 21% to 69 ± 42% lower than the NPP of a completely intact canopy (Table 2). While previous studies (Kurz et al. 2008, Clark et al. 2010, Schäfer et al. 2010, Heliasz et al. 2011, Metcalfe et al. 2014, Lund et al. 2017) have shown that herbivore outbreaks can result in a considerable loss of carbon from the ecosystem, our study suggests that insect herbivores can have a large impact on the forest primary productivity even at a low density.

**Table 2.** Oak net primary productivity (NPP) for intact canopy and canopy under normal level of herbivory and the effect of herbivory on the NPP estimated with different methods (canopy upscaling, biometric measurements, and tree census) and at different scales: a hypothetical plot consisting only of oak trees (basal area of 33.3 m^2^ ha^−1^), and the measured site (basal area of oaks 6.7 m^2^ ha^−1^, 20% of the total basal area). The errors are propagated measuring errors and uncertainties related to the different scaling functions. The unit in the table is Mg C ha^−1^ year^−1^.

|  | **NPP**<br>**Normal**<br>**herbivory** | **±**<br>**error** | **NPP**<br>**Intact**<br>**canopy** | **±**<br>**error** | **Herbivory**<br>**effect, C** | **±**<br>**error** | **Herbivory**<br>**effect %** | **±**<br>**error** |
| --- | --- | --- | --- | --- | --- | --- | --- | --- |
| **Per plot of only oak** |  |  |  |  |  |  |  |  |
| Canopy upscaling | 3.8 | 4.9 | 12.3 | 8.2 | 8.5 | 5.1 | 69.4 | 41.6 |
| Biometric measurements | 4.6 | 1.1 | 10.0 | 1.8 | 5.4 | 2.1 | 54.3 | 20.9 |
| Tree census | 3.5 | 0.6 | 7.5 | 2.2 | 4.1 | 2.3 | 54.3 | 30.4 |
| **Per real plot (counting only oak trees)** |  |  |  |  |  |  |  |  |
| Canopy upscaling | 0.8 | 1.0 | 2.5 | 1.7 | 1.7 | 1.0 | 69.4 | 41.6 |
| Biometric measurements | 0.9 | 0.2 | 2.0 | 0.4 | 1.1 | 0.4 | 54.3 | 20.9 |
| Tree census | 0.6 | 0.1 | 1.2 | 0.4 | 0.7 | 0.4 | 54.3 | 30.4 |

**Figure 3.**
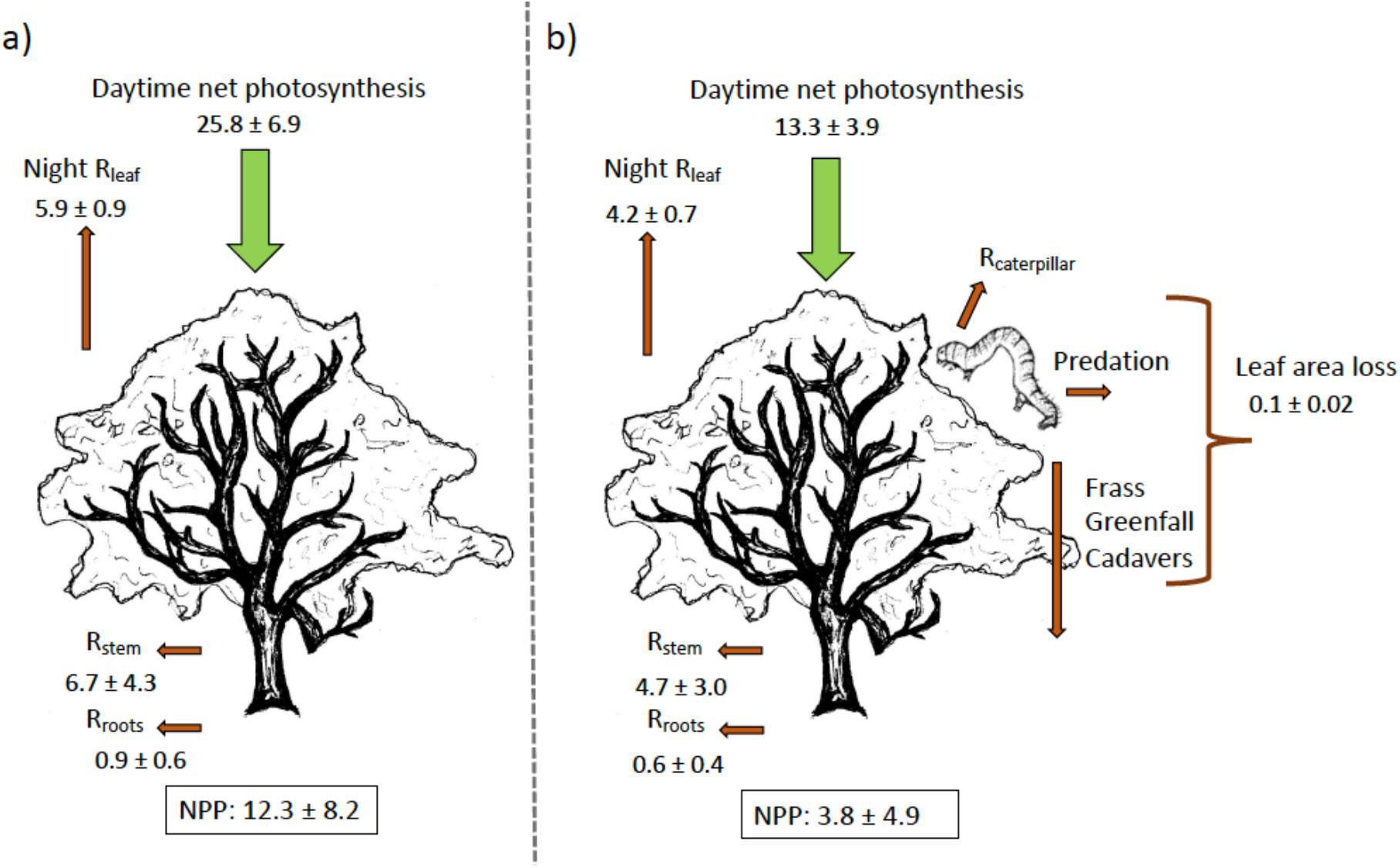
The combined carbon budget a) without and b) with herbivory for an oak forest as Mg C ha^−1^ yr^−1^ based on canopy upscaling. The carbon flux of leaf area loss (0.1 Mg C ha^−1^ yr^−1^) will continue its cycle either by entering soil as frass or insect cadavers, being respired by herbivores, or being eaten by higher trophic levels preying on caterpillars (see Figure 5). Errors are propagated measuring uncertainties. R - respiration, NPP – net primary productivity (= Photosynthesis – (R_leaf_ + R_stem_ + R_roots_)).

We suggest that the relationship between the intensity of herbivory and its effect on ecosystem productivity is non-linear and described by two stages: first, a rapid, linear increase in the effect of herbivory due to the spread of systemic effects in the canopy (suppression of photosynthesis), and second, a slower non-linear increase with increasing leaf area loss (Figure 4). Starting with a completely intact canopy, as the level of herbivory increases, the proportion of intact, unaffected leaves decreases until the level of ca. 5% herbivory (Appendix 2, Figure S6a). At this point, all shoots are likely to have at least one damaged leaf, and consequently all leaves would be affected by herbivory (either directly or through systemic effects). After this point, further changes in canopy photosynthesis are caused entirely by the direct effects of leaf area loss (Appendix 2, Figure S6cd). Assuming that the photosynthetic rate of intact leaves increases with herbivory for example as a compensatory response (Thomson et al. 2003, Retuerto et al. 2004), or that part of the missing leaf area is compensated through a second leaf flush reduces the magnitude of these estimates, but not the shape of the relationship (Appendix 3).

**Figure 4.**
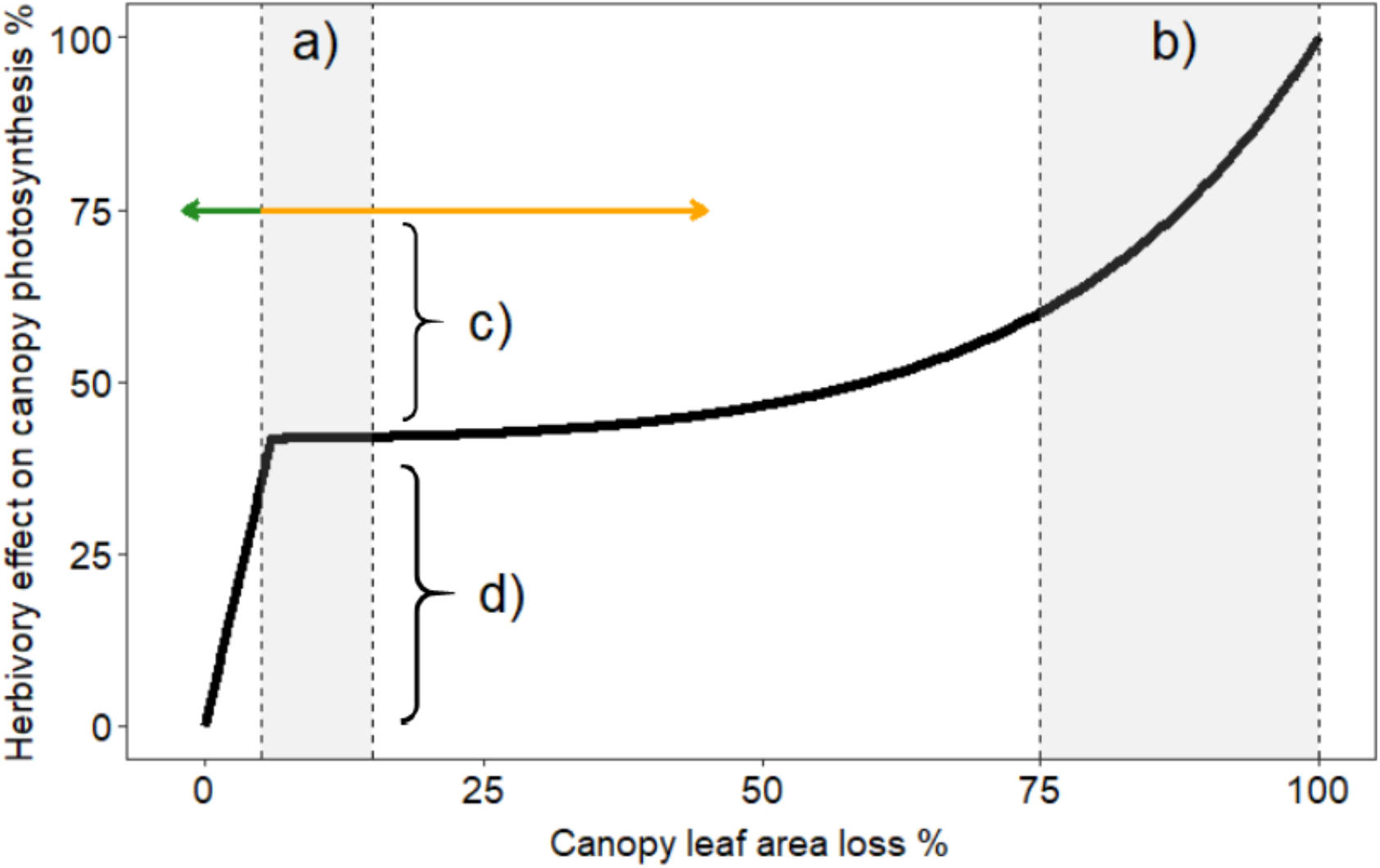
The effect of leaf area loss on daytime canopy net photosynthesis, as % reduction from the full photosynthetic capacity. The first shaded area (a) represents a typical level of herbivory in temperate forests (5-15% of leaf area loss; (Cebrian 1999). The second shaded area (b) shows leaf area loss during an insect herbivore outbreak (75-100% leaf area loss, e.g. Clark et al. 2010, Schäfer et al. 2010). At low herbivory levels, below the level at which all leaves are affected by herbivory, the herbivory effect on photosynthesis is mainly driven by the spread of systemic changes in leaf gas exchange in the undamaged leaf tissue (green arrow, “indirect effects”). Above this level, the changes in the effect of herbivory are driven by increasing leaf area loss (orange arrow). Most previous studies on the effects of herbivory on primary productivity have compared outbreak densities of herbivores to a normal level of herbivory (c). Nevertheless, we predict this effect might be smaller than comparing normal levels of herbivory to a completely intact canopy (d). See Appendix 2, Figure S6 for description of the changes in the proportion of different leaf types and leaf area loss with changing herbivore densities and Appendix 3 for relationships between insect herbivory and its effect on canopy assimilation with different underlying assumptions.

Our study illustrates the potential magnitude of the effect of insect herbivores on forest carbon cycling. Nevertheless, our estimates are based on the upscaling of leaf-level measurements and on allometric relationships, rather than on direct measurements of canopies and forest stands with and without herbivory. Below, we compare our results to previous estimates of the effects of herbivores on forest NPP. We discuss the limitations of our models and the discrepancies between our simulation and the real-world effects of herbivores on ecosystem productivity.

### Previous estimates of the effects of insect herbivores on carbon cycling

The estimates of oak forest NPP under ambient herbivory levels derived through the three different methods (canopy upscaling, tree census and biometric measurements; Table 2) were similar to one another, and agree with previous estimates of NPP at the site (Fenn et al. 2015) and at other oak-dominated temperate forests (e.g. Whitehead et al. 2004; Schäfer et al. 2010). The observed leaf area loss was 5.9 %, which is close to the reported global average for forests (5%; Cebrian 1999, 2004). We estimated that canopy gas exchange was reduced by 54%, and whole tree NPP by 54-69% due to herbivory, depending on the estimation method. These values are similar to previously published results: based on observed changes in sap flow and canopy modelling, Schäfer et al. (2010) estimated that a gypsy moth outbreak reduced canopy assimilation of an oak/pine forest by 24%. Focusing on the same outbreak, Clark et al. (2010) estimated on average 55% reduction in net ecosystem CO_2_ exchange using eddy covariance and biometric measurements (Table 3). In line with our simulations, (Flower and Gonzalez-Meler 2015) showed in a meta-analysis how the relationship between pest outbreak intensity and its effect on forest NPP was sigmoidal rather than linear, though the effects of pests were evident only after at least 30% of the basal area was affected. Comparing an outbreak year to a typical (non-outbreak) level of herbivory is likely to yield different results from the kind of comparison made in our study: even in years of low herbivore density, forest canopies experience some herbivory (Cebrian 1999, 2004). Since the indirect suppression of photosynthesis in intact plant tissue was the biggest driver of the effect of herbivory on NPP, comparing a plant experiencing a low level of herbivory to a completely intact plant might reveal a larger effect of herbivory than comparing a normal level of herbivory to an outbreak level (Figure 4).

**Table 3.** Previous estimates on the effects of insect herbivores on ecosystem carbon cycling. The baseline estimate represents productivity under normal level of herbivory (e.g. before an insect outbreak), or in the absence of herbivores (this study and Dungan et al. 2007). Herbivory level is percentage of leaf area loss, unless otherwise stated.

| Reference | Baseline<br>Mg C<br>ha<sup>-1</sup> yr<sup>-1</sup> | Baseline<br>measure | Herbivore<br>effect Mg<br>C ha<sup>-1</sup> yr<sup>-1</sup> | Herbivore<br>effect % | Method | Study system | Herbivore species | Herbivory<br>level |
| --- | --- | --- | --- | --- | --- | --- | --- | --- |
| Nardini et al. 2004 | 5 | NPP | -4.3 | -85% | Gas exchange<br>and canopy<br>model | Horse chestnut<br>(*Aesculus*<br>*hippocastanum*) | Leafminer<br>(*Cameraria ohridella*) | 25-60% of<br>LAI |
| Dungan et al. 2007 | 17.2 | Canopy<br>photosynthesis | 0.7 | 4% | Canopy model | Beech<br>(*Nothofagus*<br>*solandri*) | Scale insect<br>(*Ultracoelostoma*<br>*assimile*) | 0 (phloem<br>feeding) |
| Cook a et al. 2008 | 3.7 | NEP | -3.0 | -79% | Model and<br>eddy<br>covariance | Temperate<br>deciduous<br>forest | Tent caterpillar<br>(*Malacosoma*<br>*disstria*) | 34% |
| Kurz et al. 2008 | - | NBP | -0.4 | - | Forest<br>ecosystem<br>model | Pine /spruce | Pine beetle<br>(*Dendroctonus*<br>*ponderosae*) | - |
| Clark et al. 2010 | 1.9 | NEP | -0.7-4.8 | [-34%, -57%] | Eddy<br>covariance<br>and biometric<br>measurements | Oak /pine | Gypsy moth<br>(*Lymantria dispar*) | > 75% |
| Schäfer et al. 2010 | 12.4 | Canopy<br>photosynthesis | -3 | -24% | Sap-flow<br>based canopy<br>modelling | Oak /pine | Gypsy moth<br>(*Lymantria dispar*) | > 75% |
| Heliasz et al. 2011 | 1.1 | NEP | -0.9 | -89% | Eddy<br>covariance | Mountain birch<br>(*Betula*<br>*pubescens*) | Autumnal moth<br>(*Epirrita autumnata*) | 100% |
| Malhi et al. 2014 | 14.2-15 | NPP | -1.5 | -11% | Estimation of<br>leaf area loss | Tropical forest | Several | 18.80% |
| Flower et al. 2012 | 3.4 | NPP | -1.5 | -43% | Biometric<br>measurements | Ash (*Fraxinus*<br>*americana*) | Emerald ash borer | - |
| Metcalfe et al. 2013 | 2-5 | Foliar production | -0.3-0.9 | [-12%, -19%] | Estimation of leaf area loss | Tropical forest | Several | 19-20% |
| Strickland et al. 2013 | - | - | - | -33% | <sup>13</sup>C pulse chase experiment | Meadow | Grasshopper (*Melanoplus femurrubrum*) | 32gm<sup>-2</sup> of biomass |
| Lund et al. 2017 | 4.5 | NEP | -2.6 | -58% | Chamber-based CO<sub>2</sub> exchange | Tundra | Moth (*Eurois occulta*) | 100% |
| **this study** | **11.1** | **Oak NPP** | **-8.5** | **-69%** | **Leaf gas exchange** | **Oak (*Quercus robur*)** | **Early-season moth caterpillars** | **6%** |
|  |  |  | **-0.1** |  | **Leaf area loss** |  |  |  |

NPP – net primary productivity, NEP – net ecosystem productivity, NBP – net biome productivity

In our study, the largest effect of herbivory on NPP (69%) was estimated through the canopy upscaling approach. This is most likely because stem and root respiration were assumed to respond to herbivory similarly to leaf respiration, and because the difference between the two scenarios (intact and with herbivory) was smaller for leaf respiration than for photosynthesis. One of the biggest uncertainties in our calculations is how woody respiration responds to leaf herbivory. This is a clear knowledge gap that needs to be addressed in future studies. Had we assumed that a change in photosynthesis would not result in a change in stem respiration, or that stem respiration would increase after herbivory, our estimate for the effect of herbivory on oak NPP would have been even larger. Thus, our current assumptions on the response of woody respiration to herbivory provide a more conservative estimate on the effect of herbivory on forest NPP than if we had assumed constant stem respiration or an increased respiration after herbivory.

### The effect of increasing herbivory pressure on forest productivity

Our estimate of the effect of herbivores on oak NPP is based on data from a year with a low winter moth population density. Previous reports of winter moth outbreaks in Wytham Woods include records of complete defoliation of trees (Feeny 1970). Intuitively, we might expect increasing herbivory pressure to have a linearly increasing effect on ecosystem productivity. Nevertheless, the herbivore-induced changes in leaf gas exchange per unit leaf area of the remaining leaf tissue appear not to depend on the amount of damage per leaf (Visakorpi et al. 2018), and the vast majority of leaves show some signs of feeding damage even under low herbivore densities. Thus, increased herbivore densities would likely not further affect the leaf gas exchange in the remaining tissue. Consequently, we predict that increases in herbivore densities above those observed in our study are unlikely to result in large changes in the magnitude of the effect of herbivory on ecosystem NPP, except in outbreak years with very high proportion of leaf area loss (e.g. 75% or more, Figure 4). If the systemic plant responses observed in our system are widespread, many forests could be at a stage where a large part of the photosynthesizing plant tissue is already indirectly affected by herbivory. Consequently, observing large-scale herbivore-induced changes in ecosystem productivity in natural conditions might be restricted to situations with insect outbreaks.

The predicted magnitude of reduction in productivity caused by reduced photosynthesis is large, and we recognise that this prediction needs explicit testing. There are several additional factors that might change how herbivory affects ecosystem productivity which have not been considered in our predictions. First, the reduced leaf area caused by herbivory could increase light penetration to lower canopy layers (Anten and Ackerly 2001) and increase canopy light use efficiency (Gough et al. 2013). The effect of increased light penetration might compensate for some of the loss of leaf tissue (e.g. 5-30% according to Anten and Ackerly, 2001). Even though our assimilation model accounts for light diffusion through the canopy, and our simulations assume that LAI changes with herbivory (Appendix 2, Figure S7), a more detailed canopy model taking into account sunlit and shaded areas could simulate the effects of increased light more accurately (Clark et al. 2011). Our model also assumes that efficiency at which canopy intercepts light (i.e. the light extinction coefficient, see Equation 1 in Appendix 1) is constant, which is most likely unrealistic. Furthermore, how herbivory is distributed between canopy layers is currently unknown, but could affect how much light reaches the lower leaves. Estimating light interception in relation to herbivory-induced changes in LAI and patterns of herbivory within canopies would allow better quantification of the effects of increased light at lower canopy layers.

Second, increased herbivory is likely to result in increased deposits of frass, insect tissue, leaf fragments (‘greenfall’) and nutrient leaching from damaged leaves (“throughfall”). Herbivore-induced reduction in tree growth could also reduce competition experienced by the surrounding vegetation. The increased nutrient cycling and competitive release could increase forest productivity and compensate for the loss of carbon to herbivory (Gough et al. 2013, Lund et al. 2017, Costilow et al. 2017). Third, increased leaf herbivory might increase carbon allocation to roots (Dyer et al. 1991), which might increase root productivity and root and soil respiration (Holland et al. 1996). Fourth, the total litterfall might remain unchanged even after severe defoliation due to compensating second leaf flushes (Grace 1986, Clark et al. 2010). If the second leaf flush (“lammas shoots” on oak) produces more leaves as a response to early-season herbivory, or if these new leaves photosynthesize at a higher rate, they might compensate for the effects of herbivory on the canopy gas exchange. Lastly, insect feeding can increase transport of photosynthetic end-products away from certain leaves, and thereby reduce the negative effects of herbivory on photosynthesis (Retuerto et al. 2004, Schwachtje and Baldwin 2008). These types of sink-source dynamics might change the distribution of photosynthetic efficiency within the canopy. The higher photosynthetic rate of intact leaves could thus be a compensatory response to herbivory (see e.g. Thomson et al. 2003, Retuerto et al. 2004).

We simulated the relationship between the effect of herbivory on canopy assimilation and the intensity of herbivory assuming compensatory photosynthesis and a second leaf flush. If photosynthesis of intact leaves increases with increasing level of herbivory (which could be assumed if the response was compensatory) the effect of herbivory on canopy assimilation would be lower than without the compensatory response, especially at low herbivore densities (e.g. 23% compared to 49% with 6% herbivory, Figure S8b in Appendix 3). A second leaf flush occurring 1^st^ of August, compensating 50% of the leaf area loss, would on the other hand lower the estimate on the effect of herbivory at high herbivore densities: the effect of herbivory would never exceed 75% (Figure S8d in Appendix 3). Compensatory plant responses are likely to be important in reducing the impact of herbivory on forest productivity, but their exact effect depends on the compensating mechanism. Quantifying the compensatory plant responses to herbivory and the photosynthetic rate of lammas leaves would be important topics for future studies in this system.

### The effect of decreasing herbivory pressure on forest productivity

Our results indicate that the lower the herbivory rate, the more important the indirect effects of herbivory on photosynthesis. When only a small part of the canopy is affected by herbivores, even small increases in herbivory could cause a large change in the canopy photosynthesis through the disproportionate decrease in the number of intact leaves (herbivory on a single leaf will trigger systemic effects on all leaves within that shoot). For example, in a study by (Zvereva et al. 2012), a small increase in low intensity herbivory (from 1% to 3% leaf area loss) in mountain birch (*Betula pubescens* subsp. *czerepanovii*) resulted in a much larger (30%) reduction in plant growth. Nevertheless, since our study represents a year with a relatively low herbivore density, situations of even lower levels of herbivory and the rapid changes in productivity might be rare in natural conditions.

Predicting the effects of reduced levels of early-season herbivory on forest carbon cycling is not straightforward. Reduced densities of herbivores early in the season could result in altered investments in plant defences and change the level of herbivory experienced by the host plant at a later point (Agrawal 2000, Poelman et al. 2008). Changes in the frequency of insect herbivory could also change the susceptibility to plant pathogens (Felton and Korth 2000, De Vos et al. 2006), which could decrease leaf photosynthesis (Copolovici et al. 2014). How the interplay between different plant enemies affects carbon sequestration and cycling is currently unknown.

The seasonal changes in oak photosynthesis in our study system (Morecroft et al. 2003) might depend on early-season herbivore densities. The slow onset and the late peak of oak photosynthesis could be a strategy to avoid losing photosynthesised sugars to early-season caterpillars and could be driven by phenotypic plasticity in the responses to the intensity of early-season herbivores. The oak photosynthesis could develop faster when the spring-feeding caterpillars occur at low density compared to a high-density year, changing the amount of carbon sequestrated over the season.

### The fate of carbon contained in the leaf area lost to herbivory

Previous studies on the patterns of leaf area loss and insect energetics allows us to estimate the pathways of carbon once it is removed from the canopy. Based on previous studies on the amounts of greenfall (falling leaf fragments due to herbivore feeding), roughly 75% of the lost leaf area is ingested by the caterpillars (Russell et al. 2004). Of the ingested carbon, approximately 15% will be respired, 60% will be turned into frass and transported into the soil, and the rest will turn into insect tissue (of lepidopteran caterpillars; Wiegert and Petersen 1983). Based on estimated predation rates of winter moth caterpillars and cadavers, most of this insect tissue will be eaten by predators (East 1974). Thus, we estimate that 18% of the carbon contained in the observed leaf area lost to herbivory (0.02 Mg C ha^−1^ yr^−1^) is likely to be transferred to higher trophic levels in this system, and 70% of the leaf area loss (0.08 Mg C ha^−1^ yr^−1^) is directly deposited in the soil, partly as greenfall and partly as frass (Figure 5). Data on the amount of plant biomass consumed by herbivores have been collected for several ecosystems (Cebrian 2004). By using similar calculations here, it could be possible to estimate herbivore-mediated carbon fluxes to soil or higher trophic levels for a wide variety of ecosystems (see e.g. Metcalfe and others 2014).

**Figure 5.**
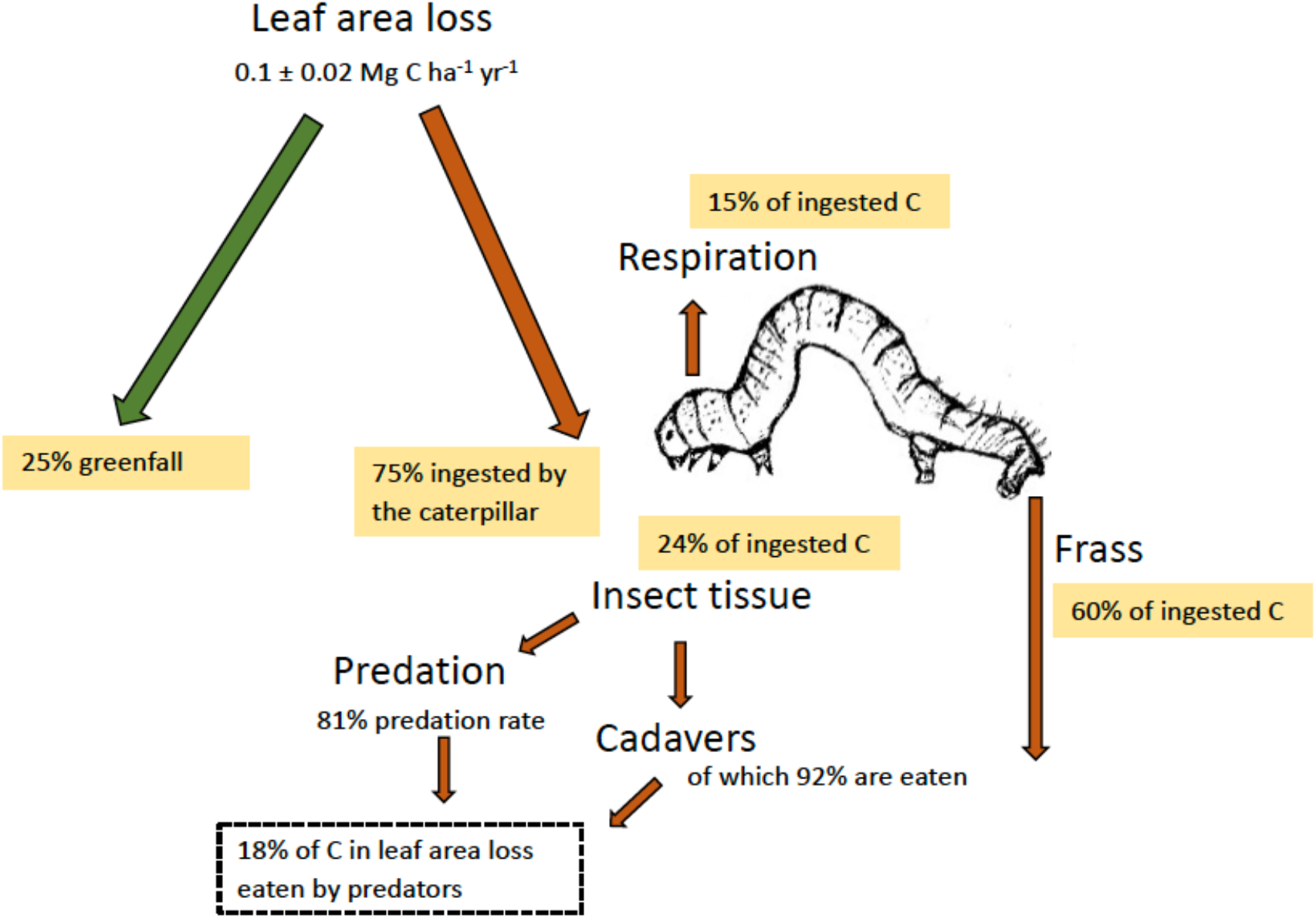
Estimating the fate of the carbon lost through eaten leaf area in the oak-caterpillar study system in Wytham Woods. Of the observed leaf area loss, roughly 75% is expected to be ingested by the caterpillar (Russell et al. 2004). Of this, 15% will be respired, 60% falls down as frass, and 24% will be used by the insect to grow and reproduce, based on earlier studies of insect energetics (Wiegert and Petersen 1983). Of winter moth caterpillars in Wytham Woods, 81% of live caterpillars and 92% of cadavers are preyed upon (East 1974). Thus, 24% of the carbon ingested by the caterpillar (or 18% of the carbon contained in the observed leaf area loss, 0.02 Mg C ha^−1^ yr^−1^), is transferred to higher trophic levels.

## Conclusions

We propose that insect herbivores can, even at low densities, have a large effect on ecosystem NPP in a temperate deciduous forest, mainly through indirect changes in the rates of canopy gas exchange. Most forest ecosystems might naturally be in a state where photosynthesis is substantially reduced by the effects of herbivory, in particular by the indirect effects. The relationship between the intensity of herbivory and its effects on the host plant is most likely non-linear. At low herbivore densities, the effect of herbivory on NPP is primarily driven by changes in leaf gas exchange in the remaining leaf tissue, while the contribution of leaf area loss increases with increasing herbivore abundances. We predict that comparisons of productivity between completely intact plants and plants with even a small amount of herbivory-inflicted damage are likely to yield large estimates on the effects of herbivory on plant productivity. On the other hand, comparisons between plants experiencing low (but non-zero) and high levels of herbivory are likely to result in lower estimates on the effect of herbivory. The presence of compensatory plant responses, like increased photosynthesis or additional leaf flushes will likely reduce the impact of herbivory, the exact effect depending on the compensating mechanism. Further and more detailed studies including different study systems and species are needed to validate whether the proposed relationship between intensity of herbivory and its effect on plant productivity holds. Where there is strong interannual variability in herbivory, ecosystem measurements of productivity, such as eddy covariance approaches (Thomas et al. 2011) or detailed estimates on tree growth (Varley and Gradwell 1962, Whittaker and Warrington 1985, Zvereva et al. 2012) could be coupled with herbivory measurements to test this relationship. The distributions and abundances of insect herbivores are likely to change in the future (Ayres and Lombardero 2000, Jepsen et al. 2008, Kurz et al. 2008), with increasing temperature (Bale et al. 2002), CO_2_ concentration (Stiling et al. 2009), drought (Gaylord et al. 2013) and advancing phenology (Charmantier et al. 2008) affecting the intensity of herbivory differently. Quantifying the effect of insect herbivores carbon cycling is therefore an intriguing and important avenue of research.

## Acknowledgements

We thank Lionel Cole for providing data on winter moth abundances, Alexander Shenkin and Kieran Walker for the stem respiration data and the Journal Club of the Ecosystems lab and three anonymous reviewers for helpful comments on the text. KV was funded by Osk.

Huttunen Foundation and Finnish Cultural Foundation. SG is a Royal Society University Research Fellow. YM was supported by the Jackson Foundation and by a European Research Council Advanced Investigator Grant (GEM-TRAIT: 321131).

## References

Agrawal, A. A. 2000. Specificity of induced resistance in wild radish: causes and consequences for two specialist and two generalist caterpillars. Oikos 89:493–500.

Amiro, B. D., A. G. Barr, J. G. Barr, T. A. Black, R. Bracho, M. Brown, J. Chen, K. L. Clark, K. J. Davis, A. R. Desai, S. Dore, V. Engel, J. D. Fuentes, A. H. Goldstein, M. L. Goulden, T. E. Kolb, M. B. Lavigne, B. E. Law, H. A. Margolis, T. Martin, J. H. McCaughey, L. Misson, M. Montes-Helu, A. Noormets, J. T. Randerson, G. Starr, and J. Xiao. 2010. Ecosystem carbon dioxide fluxes after disturbance in forests of North America. Journal of Geophysical Research 115:G00K02.

Anten, N. P. R., and D. D. Ackerly. 2001. Canopy-level photosynthetic compensation after defoliation in a tropical understorey palm. Functional Ecology 15.

Ayres, M. P., and M. J. Lombardero. 2000. Assessing the consequences of global change for forest disturbance from herbivores and pathogens. Science of The Total Environment 262:263–286.

Bale, J. S., G. J. Masters, I. D. Hodkinson, C. Awmack, T. M. Bezemer, V. K. Brown, J. Butterfield, A. Buse, J. C. Coulson, J. Farrar, J. E. G. Good, R. Harrington, S. Hartley, T. H. Jones, R. L. Lindroth, M. C. Press, I. Symrnioudis, A. D. Watt, and J. B. Whittaker. 2002. Herbivory in global climate change research: direct effects of rising temperature on insect herbivores. Global Change Biology 8:1–16.

Bilgin, D. D., J. A. Zavala, J. Zhu, S. J. Clough, D. R. Ort, and E. H. DeLucia. 2010. Biotic stress globally downregulates photosynthesis genes. Plant, Cell & Environment 33:1597–1613.

Butt, N., G. Campbell, Y. Malhi, M. Morecroft, K. Fenn, and M. Thomas. 2009. Initial results from establishment of a long-term broadleaf monitoring plot at Wytham Woods, Oxford, UK. University of Oxford, Oxford.

Campioli, M., Y. Malhi, S. Vicca, S. Luyssaert, D. Papale, J. Peñuelas, M. Reichstein, M. Migliavacca, M. A. Arain, and I. A. Janssens. 2016. Evaluating the convergence between eddy-covariance and biometric methods for assessing carbon budgets of forests. Nature Communications 7:13717.

Cebrian, J. 1999. Patterns in the fate of production in plant communities. The American Naturalist 154:449–468.

Cebrian, J. 2004. Role of first-order consumers in ecosystem carbon flow. Ecology Letters 7:232–240.

Chapin, F. S. 1997. Biotic Control over the Functioning of Ecosystems. Science 277:500–504.

Charmantier, A., R. H. McCleery, L. R. Cole, C. Perrins, L. E. B. Kruuk, and B. C. Sheldon. 2008. Adaptive phenotypic plasticity in response to climate change in a wild bird population. Science 320:800–803.

Clark, D. B., L. M. Mercado, S. Sitch, C. D. Jones, N. Gedney, M. J. Best, M. Pryor, G. G. Rooney, R. L. H. Essery, E. Blyth, O. Boucher, R. J. Harding, C. Huntingford, and P. M. Cox. 2011. The Joint UK Land Environment Simulator (JULES), model description – Part 2: Carbon fluxes and vegetation dynamics. Geoscientific Model Development 4:701–722.

Clark, K. L., N. Skowronski, and J. Hom. 2010. Invasive insects impact forest carbon dynamics: defoliation and forest carbon dynamics. Global Change Biology 16:88–101.

Copolovici, L., F. Vaartnou, M. P. Estrada, and U. Niinemets. 2014. Oak powdery mildew (Erysiphe alphitoides)-induced volatile emissions scale with the degree of infection in Quercus robur. Tree Physiology 34:1399–1410.

Costilow, K. C., K. S. Knight, and C. E. Flower. 2017. Disturbance severity and canopy position control the radial growth response of maple trees (Acer spp.) in forests of northwest Ohio impacted by emerald ash borer (Agrilus planipennis). Annals of Forest Science 74:10.

De Vos, M., W. van Zaalen, A. Koornneef, J. P. Korzelius, M. Dicke, L. C. Van Loon, and C. M. J. Pieterse. 2006. Herbivore-induced resistance against microbial pathogens in Arabidopsis. Plant Physiology 142:352–363.

Dyer, M. I., M. A. Acra, G. M. Wang, D. C. Coleman, D. W. Freckman, S. J. McNaughton, and B. R. Strain. 1991. Source-sink carbon relations in two Panicum coloratum ecotypes in response to herbivory. Ecology 72:1472–1483.

East, R. 1974. Predation on the soil-dwelling stages of the winter moth at Wytham Woods, Berkshire. The Journal of Animal Ecology 43:611.

Edburg, S. L., J. A. Hicke, P. D. Brooks, E. G. Pendall, B. E. Ewers, U. Norton, D. Gochis, E. D. Gutmann, and A. J. Meddens. 2012. Cascading impacts of bark beetle-caused tree mortality on coupled biogeophysical and biogeochemical processes. Frontiers in Ecology and the Environment 10:416–424.

Estes, J. A., J. Terborgh, J. S. Brashares, M. E. Power, J. Berger, W. J. Bond, S. R. Carpenter, T. E. Essington, R. D. C. Holt, J. B. Jackson, R. J. Marquis, L. Oksanen, T. Oksanen, R. T. Paine, E. K. Pikitch, W. J. Ripple, S. A. Sandin, M. Scheffer, T. W. Schoener, J. B. Shurin, A. R. E. Sinclair, M. E. Soulé, R. Virtanen, and D. A. Wardle. 2011. Trophic downgrading of planet Earth. Science 333:301–306.

Feeny, P. 1970. Seasonal changes in oak leaf tannins and nutrients as a cause of spring feeding by Winter moth caterpillars. Ecology 51:565–581.

Felton, G. W., and K. L. Korth. 2000. Trade-offs between pathogen and herbivore resistance. Current Opinion in Plant Biology 3:309–314.

Fenn, K. M. 2010. Carbon cycling in British deciduous woodland: processes, budgets, climate & phenology. DPhil, University of Oxford.

Fenn, K., Y. Malhi, M. Morecroft, C. Lloyd, and M. Thomas. 2015. The carbon cycle of a maritime ancient temperate broadleaved woodland at seasonal and annual scales. Ecosystems 18:1–15.

Flower, C. E., and M. A. Gonzalez-Meler. 2015. Responses of Temperate Forest Productivity to Insect and Pathogen Disturbances. Annual Review of Plant Biology 66:547–569.

Flower, C. E., K. S. Knight, and M. A. Gonzalez-Meler. 2013. Impacts of the emerald ash borer (Agrilus planipennis Fairmaire) induced ash (Fraxinus spp.) mortality on forest carbon cycling and successional dynamics in the eastern United States. Biological Invasions 15:931–944.

Forister, M. L., V. Novotny, A. K. Panorska, L. Baje, Y. Basset, P. T. Butterill, L. Cizek, P. D. Coley, F. Dem, I. R. Diniz, P. Drozd, M. Fox, A. E. Glassmire, R. Hazen, J. Hrcek, J. P. Jahner, O. Kaman, T. J. Kozubowski, T. A. Kursar, O. T. Lewis, J. Lill, R. J. Marquis, S. E. Miller, H. C. Morais, M. Murakami, H. Nickel, N. A. Pardikes, R. E. Ricklefs, M. S. Singer, A. M. Smilanich, J. O. Stireman, S. Villamarín-Cortez, S. Vodka, M. Volf, D. L. Wagner, T. Walla, G. D. Weiblen, and L. A. Dyer. 2015. The global distribution of diet breadth in insect herbivores. Proceedings of the National Academy of Sciences 112:442–447.

Gaylord, M. L., T. E. Kolb, W. T. Pockman, J. A. Plaut, E. A. Yepez, A. K. Macalady, R. E. Pangle, and N. G. McDowell. 2013. Drought predisposes piñon-juniper woodlands to insect attacks and mortality. New Phytologist 198:567–578.

Gough, C. M., B. S. Hardiman, L. E. Nave, G. Bohrer, K. D. Maurer, C. S. Vogel, K. J. Nadelhoffer, and P. S. Curtis. 2013. Sustained carbon uptake and storage following moderate disturbance in a Great Lakes forest. Ecological Applications 23:1202–1215.

Grace, J. R. 1986. The influence of gypsy moth on the composition and nutrient content of litter fall in a pennsylvania oak forest. Forest Science 32:855–870.

Griffin, K. L., D. T. Tissue, M. H. Turnbull, W. Schuster, and D. Whitehead. 2001. Leaf dark respiration as a function of canopy position in Nothofagus fusca trees grown at ambient and elevated CO2 partial pressures for 5 years. Functional Ecology 15:497–505.

Heliasz, M., T. Johansson, A. Lindroth, M. Mölder, M. Mastepanov, T. Friborg, T. V. Callaghan, and T. R. Christensen. 2011. Quantification of C uptake in subarctic birch forest after setback by an extreme insect outbreak: carbon uptake setback by insect outbreak. Geophysical Research Letters 38.

Holland, J. N., W. Cheng, and D. A. Crossley. 1996. Herbivore-induced changes in plant carbon allocation: assessment of below-ground C fluxes using carbon-14. Oecologia 107:87–94.

Hunter, M. D. 1992. Interactions within herbivore communities mediated by the host plant: the keystone herbivore concept. Pages 287–325 *in* M. D. Hunter, T. Ohgushi, and P. W. Price, editors. Effects of resource distribution on animal–plant interactions. Academin Press, Inc., San Diego, California, USA.

Jepsen, J. U., S. B. Hagen, R. A. Ims, and N. G. Yoccoz. 2008. Climate change and outbreaks of the geometrids Operophtera brumata and Epirrita autumnata in subarctic birch forest: evidence of a recent outbreak range expansion. Journal of Animal Ecology 77:257–264.

Kurz, W. A., C. C. Dymond, G. Stinson, G. J. Rampley, E. T. Neilson, A. L. Carroll, T. Ebata, and L. Safranyik. 2008. Mountain pine beetle and forest carbon feedback to climate change. Nature 452:987–990.

Lund, M., K. Raundrup, A. Westergaard-Nielsen, E. López-Blanco, J. Nymand, and P. Aastrup. 2017. Larval outbreaks in West Greenland: Instant and subsequent effects on tundra ecosystem productivity and CO2 exchange. Ambio 46:26–38.

Mercado, L. M., C. Huntingford, J. H. C. Gash, P. M. Cox, and V. Jogireddy. 2007. Improving the representation of radiation interception and photosynthesis for climate model applications. Tellus B: Chemical and Physical Meteorology 59:553–565.

Metcalfe, D. B., G. P. Asner, R. E. Martin, J. E. Silva Espejo, W. H. Huasco, F. F. Farfán Amézquita, L. Carranza-Jimenez, D. F. Galiano Cabrera, L. D. Baca, F. Sinca, L. P. Huaraca Quispe, I. A. Taype, L. E. Mora, A. R. Dávila, M. M. Solórzano, B. L. Puma Vilca, J. M. Laupa Román, P. C. Guerra Bustios, N. S. Revilla, R. Tupayachi, C. A. J. Girardin, C. E. Doughty, and Y. Malhi. 2014. Herbivory makes major contributions to ecosystem carbon and nutrient cycling in tropical forests. Ecology Letters 17:324–332.

Meza-Canales, I. D., S. Meldau, J. A. Zavala, and I. T. Baldwin. 2017. Herbivore perception decreases photosynthetic carbon assimilation and reduces stomatal conductance by engaging 12-oxo-phytodienoic acid, mitogen-activated protein kinase 4 and cytokinin perception. Plant, Cell & Environment 40:1039–1056.

Monsi, M., and T. Saeki. 1953. Ueber den Lichtfaktor in den Planzengesellschaften und seine Bedeutung fuer die Stoffproduktion. Journal of Japanese Botany:22–52.

Morecroft, M. D., V. J. Stokes, and J. I. L. Morison. 2003. Seasonal changes in the photosynthetic capacity of canopy oak (Quercus robur) leaves: the impact of slow development on annual carbon uptake. International Journal of Biometeorology 47:221–226.

Nabity, P. D., J. A. Zavala, and E. H. DeLucia. 2009. Indirect suppression of photosynthesis on individual leaves by arthropod herbivory. Annals of Botany 103:655–663.

Nykänen, H., and J. Koricheva. 2004. Damage-induced changes in woody plants and their effects on insect herbivore performance: a meta-analysis. Oikos 104:247–268.

Oleksyn, J., P. Karolewski, M. J. Giertych, R. Zytkowiak, P. B. Reich, and M. G. Tjoelker. 1998. Primary and secondary host plants differ in leaf-level photosynthetic response to herbivory: evidence from Alnus and Betula grazed by the alder beetle, Agelastica alni. New Phytologist 140:239–249.

Poelman, E. H., C. Broekgaarden, J. J. A. Van Loon, and M. Dicke. 2008. Early season herbivore differentially affects plant defence responses to subsequently colonizing herbivores and their abundance in the field. Molecular Ecology 17:3352–3365.

Rennie, S., J. Adamson, R. Anderson, C. Andrews, J. Bater, N. Bayfield, K. Beaton, D. Beaumont, S. Benham, V. Bowmaker, C. Britt, R. Brooker, D. Brooks, J. Brunt, G. Common, R. Cooper, S. Corbett, N. Critchley, P. Dennis, J. Dick, B. Dodd, N. Dodd, N. Donovan, J. Easter, E. Eaton, M. Flexen, A. Gardiner, D. Hamilton, P. Hargreaves, M. Hatton-Ellis, M. Howe, J. Kahl, M. Lane, S. Langan, D. Lloyd, B. McCarney, Y. McElarney, C. McKenna, S. McMillan, F. Milne, L. Milne, M. Morecroft, M. Murphy, A. Nelson, H. Nicholson, D. Pallett, D. Parry, I. Pearce, G. Pozsgai, R. Rose, S. Schafer, T. Scott, L. Sherrin, C. Shortall, R. Smith, P. Smith, R. Tait, C. Taylor, M. Taylor, M. Thurlow, A. Turner, K. Tyson, H. Watson, M. Whittaker, M. Wilkinson, and C. Wood. 2017. UK Environmental Change Network (ECN) meteorology data: 1991-2015. NERC Environmental Information Data Centre.

Retuerto, R., B. Fernandez-Lema, Rodriguez-Roiloa, and J. R. Obeso. 2004. Increased photosynthetic performance in holly trees infested by scale insects. Functional Ecology 18:664–669.

Russell, C. A., K. R. Kosola, E. A. Paul, and G. P. Robertson. 2004. Nitrogen cycling in poplar stands defoliated by insects. Biogeochemistry 68:365–381.

Savill, P. S., editor. 2011. Wytham woods: Oxford’s ecological laboratory. Oxford Univ. Press, Oxford.

Schäfer, K. V. R., K. L. Clark, N. Skowronski, and E. P. Hamerlynck. 2010. Impact of insect defoliation on forest carbon balance as assessed with a canopy assimilation model. Global Change Biology 16:546–560.

Schmitz, O. J., P. A. Raymond, J. A. Estes, W. A. Kurz, G. W. Holtgrieve, M. E. Ritchie, D. E. Schindler, A. C. Spivak, R. W. Wilson, M. A. Bradford, V. Christensen, L. Deegan, V. Smetacek, M. J. Vanni, and C. C. Wilmers. 2014. Animating the carbon cycle. Ecosystems 17:344–359.

Schoonhoven, L. M., J. J. A. van Loon, and M. Dicke. 2005. Insect-plant biology. 2nd ed. Oxford University Press, Oxford, UK.

Schwachtje, J., and I. T. Baldwin. 2008. Why does herbivore attack reconfigure primary metabolism? Plant Physiology 146:845–851.

Stiling, P., D. Moon, A. Rossi, B. A. Hungate, and B. Drake. 2009. Seeing the forest for the trees: long-term exposure to elevated CO 2 increases some herbivore densities. Global Change Biology 15:1895–1902.

Stokes, V. 2000. Effects of microenvironment and leaf developmental characteristics on annual carbon gain and water use in two deciduous tree species. DPhil, University of Oxford.

Strickland, M. S., D. Hawlena, A. Reese, M. A. Bradford, and O. J. Schmitz. 2013. Trophic cascade alters ecosystem carbon exchange. Proceedings of the National Academy of Sciences 110:11035–11038.

Strong, D. R., J. H. Lawton, and R. Southwood. 1984. Insects on Plants. First edition. Blackwells Scientific Publications, Southampton, United Kingdom.

Thomas, M. V., Y. Malhi, K. M. Fenn, J. B. Fisher, M. D. Morecroft, C. R. Lloyd, M. E. Taylor, and D. D. McNeil. 2011. Carbon dioxide fluxes over an ancient broadleaved deciduous woodland in southern England. Biogeosciences 8:1595–1613.

Thomson, V., S. Cunnigham, M. Ball, and A. Nicotra. 2003. Compensation for herbivory by Cucumis sativus through increased photosynthetic capacity and efficiency. Oecologia 134:167–175.

Varley, G. C., and G. R. Gradwell. 1962. The effect of partial defoliation by caterpillars on the timber production by oak trees in England. Proceedings of the XIth International Congress of Entomology (Vienna 1960) 2:211–214.

Visakorpi, K., S. Gripenberg, Y. Malhi, C. Bolas, I. Oliveras, N. Harris, S. Rifai, and T. Riutta. 2018. Small-scale indirect plant responses to insect herbivory could have major impacts on canopy photosynthesis and isoprene emission. New Phytologist 220.

Visakorpi, K., T. Riutta, Y. Malhi, J.-P. Salminen, N. Salinas, and S. Gripenberg. 2020. Changes in oak (Quercus robur) photosynthesis after winter moth (Operophtera brumata) herbivory are not explained by changes in chemical or structural leaf traits. PLOS ONE 15:e0228157.

Walker, K. 2017. Variation and patterns of CO2 efflux and sapwood content of three deciduous broadleaved trees. B.S. thesis, University of Oxford, Oxford, UK.

Wertin, T. M., and R. O. Teskey. 2008. Close coupling of whole-plant respiration to net photosynthesis and carbohydrates. Tree Physiology 28:1831–1840.

Whitehead, D., K. L. Griffin, M. H. Turnbull, D. T. Tissue, V. C. Engel, K. J. Brown, W. S. Schuster, and A. S. Walcroft. 2004. Response of total night-time respiration to differences in total daily photosynthesis for leaves in a *Quercus rubra* L. canopy: implications for modelling canopy CO _2_ exchange. Global Change Biology 10:925–938.

Whittaker, J. B., and S. Warrington. 1985. An Experimental Field Study of Different Levels of Insect Herbivory Induced By Formica rufa Predation on Sycamore (Acer pseudoplatanus) III. Effects on Tree Growth. The Journal of Applied Ecology 22:797.

Wiegert, R. G., and C. E. Petersen. 1983. Energy transfer in insects. Annual Review of Entomology 28:455–486.

Wilmers, C. C., and O. J. Schmitz. 2016. Effects of gray wolf-induced trophic cascades on ecosystem carbon cycling. Ecosphere 7:e01501.

Zangerl, A. R., J. G. Hamilton, T. J. Miller, A. R. Crofts, K. Oxborough, M. R. Berenbaum, and E. H. DeLucia. 2002. Impact of folivory on photosynthesis is greater than the sum of its holes. Proceedings of the National Academy of Sciences 99:1088–1091.

Zimov, N. S., S. A. Zimov, A. E. Zimova, G. M. Zimova, V. I. Chuprynin, and F. S. Chapin. 2009. Carbon storage in permafrost and soils of the mammoth tundra-steppe biome: Role in the global carbon budget. Geophysical Research Letters 36.

Zvereva, E. L., V. Zverev, and M. V. Kozlov. 2012. Little strokes fell great oaks: minor but chronic herbivory substantially reduces birch growth. Oikos 121:2036–2043.

