## Appendix 1 for "Does insect herbivory suppress ecosystem productivity? Evidence from a temperate woodland"

### Appendix 1. Detailed methods

**Meteorological data.** To describe the meteorological conditions at our study site, and to reconstruct photosynthetic and respiration rates for the whole growing season, we used hourly temperature, rainfall and irradiance data recorded in Wytham Woods (Rennie et al. 2017). These measurements were taken from an area of semi-natural grassland at an altitude of 160 m., known as Upper Seeds (51° 46' 14.31" N, 1° 19' 56.53" W) –an open site with the nearest trees 100 m from the Automatic Weather Station (AWS). The AWS is to a standard specification adopted for the UK Environmental Change Network, using a system supplied by Didcot Instruments (now ELE International Ltd., Hemel Hempstead, UK) all of which are based around Campbell CR10 dataloggers recording hourly integrals (Logan, Utah, USA).

**Tree height and crown area.** To scale up estimates of leaf area loss to plot level and measurements of leaf respiration to canopy level, we measured tree height, canopy height and canopy area of the five studied oaks in Wytham Woods with a clinometer (TruPulse 200™ laser hypsometer) on 25$^{th}$ November 2018.

**Herbivory.** To estimate the level of insect herbivory throughout the growing season, we used earlier estimates of herbivory on the five oak trees (Visakorpi et al. 2018). These estimates were acquired by collecting 15 shoots from each of the five oaks at four time points (16 - 28$^{th}$ May, 25$^{th}$ June, 14$^{th}$ July - 10$^{th}$ August and 18$^{th}$ August 2015). The leaves were pressed and scanned, and the remaining leaf area and the area lost to herbivory were estimated using ImageJ (NIH, MD, USA) software. Because most of the leaf damage occurred at the beginning of the season, and there were no seasonal trends in the amount of leaf area loss, we used an average leaf area loss value per tree. Using the same survey data, we also estimated the proportion of three different leaf types on each tree: completely intact leaves which are growing on shoots with only other intact leaves ("completely intact"), intact leaves growing on shoots with leaves damaged by herbivores on them ("systemically affected"), and leaves damaged by herbivores ("damaged").

To estimate the amount of carbon lost through the eaten leaf tissue, we used estimates for leaf mass per area ("LMA", g/m$^2$) and leaf carbon content (% of dry mass), which were measured for the same three leaf types from the same five trees (Visakorpi et al. 2020). To calculate the leaf area loss as m$^2$, the percentage of leaf area loss per tree was multiplied with the estimated leaf area of that tree. Leaf area was estimated by multiplying the full leaf area index ("LAI") of the site (6.5 m$^2$/m$^2$, Fenn 2010) with the crown area of each tree. The estimate for leaf area loss was then multiplied with leaf mass per area, and with leaf carbon content to achieve the amount of carbon lost to herbivory due to direct leaf area loss. The measuring errors were propagated using the sampling errors in leaf area loss, LMA, leaf carbon content and the estimated measuring error (10%) of the crown area.

**Oak NPP.** We used three methods to estimate oak net primary productivity ("NPP"). First, NPP was calculated as the difference between net canopy photosynthesis and woody respiration (sum of stem and root respiration). We scaled up leaf-level photosynthesis and respiration measures to canopy-level values and combined these with estimates on oak stem and root respiration at the site obtained from previous studies ("*NPP through canopy upscaling*"). Second, we used census data on tree growth at the site to estimate woody growth, and allometric equations for oak to estimate leaf production and belowground production ("*NPP through tree growth census*"). Third, we used earlier estimates of oak above- and belowground NPP at the site ("*NPP through biometric measurements*") measured during 2007 and 2008 (Fenn 2010). For each of the three methods, we estimate NPP for two situations. First, to make our estimates comparable to previous studies on NPPs per hectare in different forest types, we calculate NPP for a hypothetical stand comprising only of mature oak trees, assigning the total basal area of the site to oaks only. Second, for a more realistic estimate of the effect of herbivory at our study site, we estimate oak NPP per ha of the actual site, where oak basal area constitutes 20% of the total basal area.

*1.) NPP through canopy upscaling*

Daytime canopy photosynthesis and respiration. To estimate daytime canopy photosynthesis and respiration, we used light-saturated photosynthetic rates measured on the five focal study trees (Visakorpi et al. 2018). In brief, we measured photosynthesis - light response curves for the five oak trees using three different leaf types (completely intact, systemically affected, damaged) in each tree. The three leaf types were a result of an experimental manipulation of herbivory, and thus represent the effect of herbivory on the physiology of a random leaf (rather than correlation between herbivore feeding preferences and photosynthetic rate). One leaf of each leaf type was measured on each tree. All measurements took place between 9am and 8pm and from 28$^{th}$ July to 7$^{th}$ August 2015.

We estimated canopy photosynthesis for each tree separately using the photosynthetic light response data and the following equation (Visakorpi et al. 2018):

$$NPC = \int_{0}^{LAI} A_{sat} * \left(\frac{PAR}{K + PAR}\right) * (e^{-k*LAI}) - (0.5 - 0.05 * \ln(PAR * e^{-k*LAI})) * R_d,$$

(Eq. 1)

where *NPC* is canopy net photosynthesis (as μmol $m^{-2}$ $s^{-1}$ of ground area), $A_{sat}$ is the light-saturated photosynthetic rate, $k$ is a light extinction coefficient, *LAI* is the canopy leaf area index, *PAR* is the light intensity at the top of the canopy and, $K$ is the light intensity at which photosynthetic rate is half of its maximum and $R_d$ is the dark respiration rate estimated from Michaelis-Menten equation.

The model estimates photosynthesis, which is reduced in lower canopy layers by reduced light availability, and daytime respiration, which is inhibited by increased light intensity ("Kok effect", (Kok 1956, Mercado et al. 2007). The light availability is assumed to be reduced according to Beer's law (Monsi and Saeki 1953). The light extinction coefficient (*k*) was set to 0.5 as a previously used estimate for broadleaf forests (Clark et al. 2011) and PAR was set to 1000 μmol $m^{-2}$ $s^{-1}$ as a standard daytime light intensity at the top of the canopy. We estimated canopy net photosynthesis for each leaf type separately, i.e. for canopies consisting of only intact, damaged or systematically affected leaves. LAI for the canopy estimate based on damaged leaves was set to 6.5 as an earlier estimate of the peak LAI of the plot (Fenn 2010). LAI for the intact canopy and the canopy consisting of systemically affected leaves was set to be 6% higher reflecting the canopy-level leaf area loss (Table 1 in the main text, Visakorpi et al., 2018). The canopy estimate based on the intact leaves only was used for the estimate of a canopy in the absence of any herbivory (hereafter "*Intact canopy*"). For estimating photosynthesis for a canopy with the observed amount of herbivory (hereafter "*Normal herbivory*"), we multiplied the estimates with the proportion of the respective leaf type observed in the canopy, and then summed these values over the three leaf types:

$$CP_{herb} = (\sum_{t=1}^{3} \mathrm{NPC_t} * \mathrm{l}_{\,t}),$$

(Eq. 2)

where $CP_{herb}$ is canopy photosynthesis under normal herbivory, *t* denotes the three different leaf types (1 = completely intact, 2 = systemically affected, 3 = damaged), $l_t$ is the proportion of leaf type *t* in the canopy. This estimate takes into account the different proportions of the three leaf types in the canopy, the respective leaf-level photosynthetic rates, the reduction in photosynthesising leaf area caused by herbivory, the effect of reduced light through the different canopy layers, and daytime respiration rate (as part of the gas exchange measurements).

Seasonality of photosynthesis. The canopy photosynthesis is seasonally affected by three phenomena. First, the leaf expansion in the spring after the budburst affects the area of photosynthesising tissue. Second, the oak photosynthetic rate per unit leaf area changes throughout the season (Morecroft et al. 2003). This change consists of a slow increase from budburst until late June, after which photosynthesis decreases slowly until late October, and then more rapidly until leaf fall (Figure S1). Third, the leaf fall in the autumn reduces the area of photosynthesising tissue.

To estimate the effect of the spring leaf expansion on canopy photosynthesis, we used measures on the sizes of expanding oak leaves in Wytham Woods during the spring of 2000 (Stokes 2000). The leaves were measured from the moment when half of the buds had burst until full leaf expansion (from 2nd May until 14th June). We measured the proportional change in leaf area over the time leaves were expanding and used these values to scale the leaf area index for that period.

To estimate reduction in LAI at any particular date in the autumn due to leaf fall, we used records of the proportion of oak leaves collected from leaf litter traps in Wytham Woods during autumn 2007 (Fenn 2010). We again calculated the proportional change in leaf area over the autumn leaf fall period and used these values to scale the leaf area index for that period. Since the autumn leaf fall was estimated only from one season, we used the average error in the change of LAI during the spring to express the uncertainty in LAI throughout the season (Table S1).

To estimate the seasonal changes in photosynthetic rate per unit leaf area, we used measurements by (Morecroft et al. 2003) in Wytham Woods for field seasons 1996, 1999 and 2000. These data consisted of photosynthetic rates (measured at 1000 μmol $m^{-2}$ $s^{-1}$ of photosynthetically active radiation, "PAR") measured at different times over the season. To estimate the proportional change in photosynthesis and LAI for each day, we built a general additive model ("GAM"), in which the proportional change in photosynthesis due to seasonal changes and reduced LAI was modelled against the day of the year. We then used this model to predict the proportional change in photosynthesis for each day of the growing season. The model predictions were further scaled so that the peak was at 100% (full LAI and full photosynthetic capacity). We then scaled our canopy photosynthesis estimates (measured during days 209 -219, 28th July - 7th August 2015) with the proportional change to achieve an estimate of the photosynthetic rate for each day in the season. These values represent instantaneous canopy photosynthetic rates per $m^2$ of ground area, when the top of the canopy is exposed to maximum daylight (1000 PAR µmol $m^{-2}$ $s^{-1}$).

We did not include any estimates of second leaf flush on oak (the "lammas" shoots), since this type of leaves has not been sampled before at our study site. Since the seasonal changes in photosynthesis include the effect of temperature, we did not apply any temperature corrections.

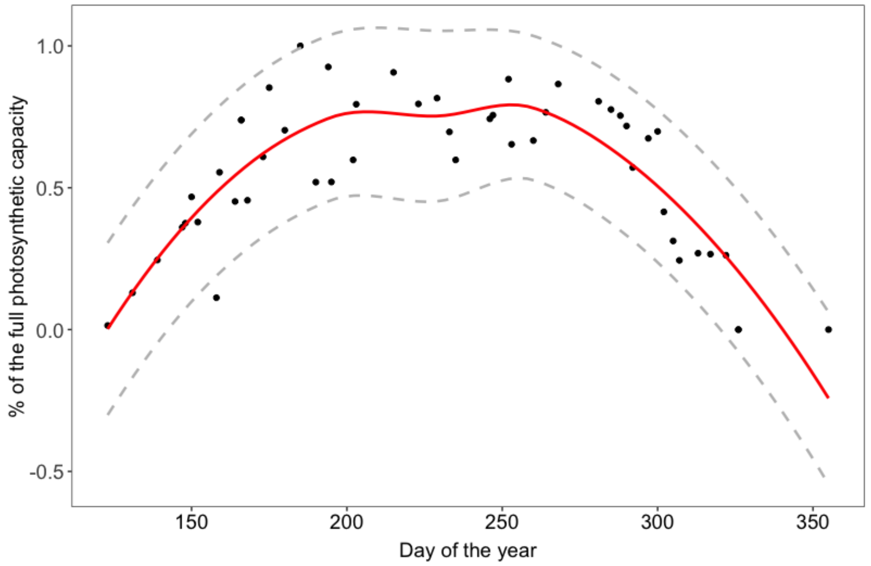


**Figure S1.** The seasonal change in photosynthetic rate per unit leaf area over the season (Morecroft et al. 2003), representing measurements over three years (1996, 1999, 2000). The dashed grey lines represent ± 1 SEM.

**Table S1.** The change in leaf length during leaf expansion in the spring (leaves reached their full size by the 14th of June), the change in leaf cover in the autumn until complete leaf fall, and the corresponding leaf area index (LAI, $m^2/m^2$). Leaf lengths are averages over sun and shade leaves and errors are ± 1 SEM (Stokes 2000). Leaf cover was estimated from litter fall traps (Fenn 2010). LAI-values were estimated by setting the full LAI to 6.5 (Fenn 2010). The LAI is lower in the spring due to smaller leaf size and in the autumn due to smaller leaf cover.

| Date | Julian date | Average leaf length (mm) | % leaf size | % leaf cover of the peak | LAI, $m^2/ m^2$ |
|---|---|---|---|---|---|
| 2nd May | 123 | 33 ± 1 | 44.0 ± 1 | - | 2.86 ± 0.03 |
| 10th May | 131 | 46 ± 4 | 61.3 ± 5 | - | 3.99 ± 0.20 |
| 18th May | 139 | 65 ± 0 | 86.7 ± 0 | - | 5.63 ± 0 |
| 26th May | 147 | 71 ± 1.5 | 94.0 ± 2 | - | 6.11 ± 0.12 |
| 31st May | 152 | 69 ± 6.5 | 91.3 ± 9 | - | 5.94 ± 0.53 |
| 14th June | 166 | 75 ± 8 | 100 | - | 6.50 ± 0 |
| 3rd Sept | 246 | - | - | 93.6 | 6.08 ± 0.15 |
| 21st Sept | 264 | - | - | 90.8 | 5.90 ± 0.15 |
| 12th Oct | 285 | - | - | 80.8 | 5.25 ± 9.15 |
| 1st Nov | 305 | - | - | 56.2 | 3.66 ± 0.15 |
| 13 Nov | 317 | - | - | 17.8 | 1.16 ± 0.15 |
| 22nd Nov | 326 | - | - | 0 | 0.00 ± 0.15 |
| 12th Dec | 355 | - | - | 0 | 0.00 ± 0.15 |

Diurnal pattern in photosynthesis. Since photosynthetic rate is strongly affected by changes in light intensity throughout the day and the season, we estimated the effect of changing daylight on photosynthetic rate for each hour for each day of the growing season (Figure S2a). We set the values calculated above as the photosynthetic rate (as μmol $CO_2$ $m^{-2}$ $s^{-1}$ of ground area) when the light intensity at the top of the canopy is 1000 μmol $m^{-2}$ $s^{-1}$ PAR. We then used hourly solar irradiation data measured 2014 (Rennie et al., 2017, see above "Meteorological data"). We used data from year 2014, because the data set for 2015 was incomplete. The irradiance was converted into PAR by multiplying the values by 4.6 (transformation from W $m^{-2}$ into photons μmol $m^{-2}$ $s^{-1}$; (Sager and McFarlane 1997) and by 0.45 (the proportion of solar irradiance at the visible light, 400-700 nm range).

To scale the canopy photosynthesis with the hourly light data, we used data on photosynthetic light response described above (Figure S2c). Since there were no differences in the proportional change in photosynthetic rate with decreasing light levels between the three different leaf types (Figure S2d), we used an average light response curve for scaling. We modelled the percentage change in photosynthesis for each observed light intensity with the Michaelis-Menten equation:

$$X_I = \frac{G_{max} I}{K + I} - R_d,$$

(Eq. 3)

where $X_I$ is the average proportional change in photosynthetic rate per light intensity *I* compared to photosynthetic rate at 1000 μmol $m^{-2}$ $s^{-1}$ of PAR (i.e. $X_{1000} = 1$), estimated from the light response curves. The maximum gross photosynthetic rate ($G_{max}$), the leaf respiration rate ($R_d$), and the light intensity at which the gross photosynthetic rate is half of its maximum (*K*, Marino et al. 2010) are model estimated parameters. We used this model to predict values for each hourly light intensity, yielding estimates for the change in photosynthesis compared to the reference level (photosynthetic rate in 1000 μmol $m^{-2}$ $s^{-1}$ PAR) for each light intensity. Since there is little evidence for mid-day water stress and stomatal closure at our study site (Morecroft and Roberts 1999), we did not include this in our estimate of the diurnal change in photosynthesis. The uncertainty in the hourly photosynthesis estimate consists of the sampling errors in the original light response curves (five point measurement per light intensity per leaf type). The errors were extrapolated for each observed hourly light intensity by building a GAM on the proportional error as the function of light intensity and extracting the fitted values of this model. These hourly estimates for the proportional change in photosynthesis and the associated error were then multiplied with the daily canopy photosynthesis estimates. This output describes instantaneous canopy photosynthetic rate as μmol $CO_2$ $m^{-2}$ $s^{-1}$ of ground area for each hour of the day. After obtaining the canopy photosynthetic rates for each hour for each day, we calculated the summed rate for each day (as "area under curve") as μmol $CO_2$ $m^{-2}$ $d^{-1}$ of ground area. We then summed these daily values to get an estimate for $CO_2$ assimilated over the whole growing season.

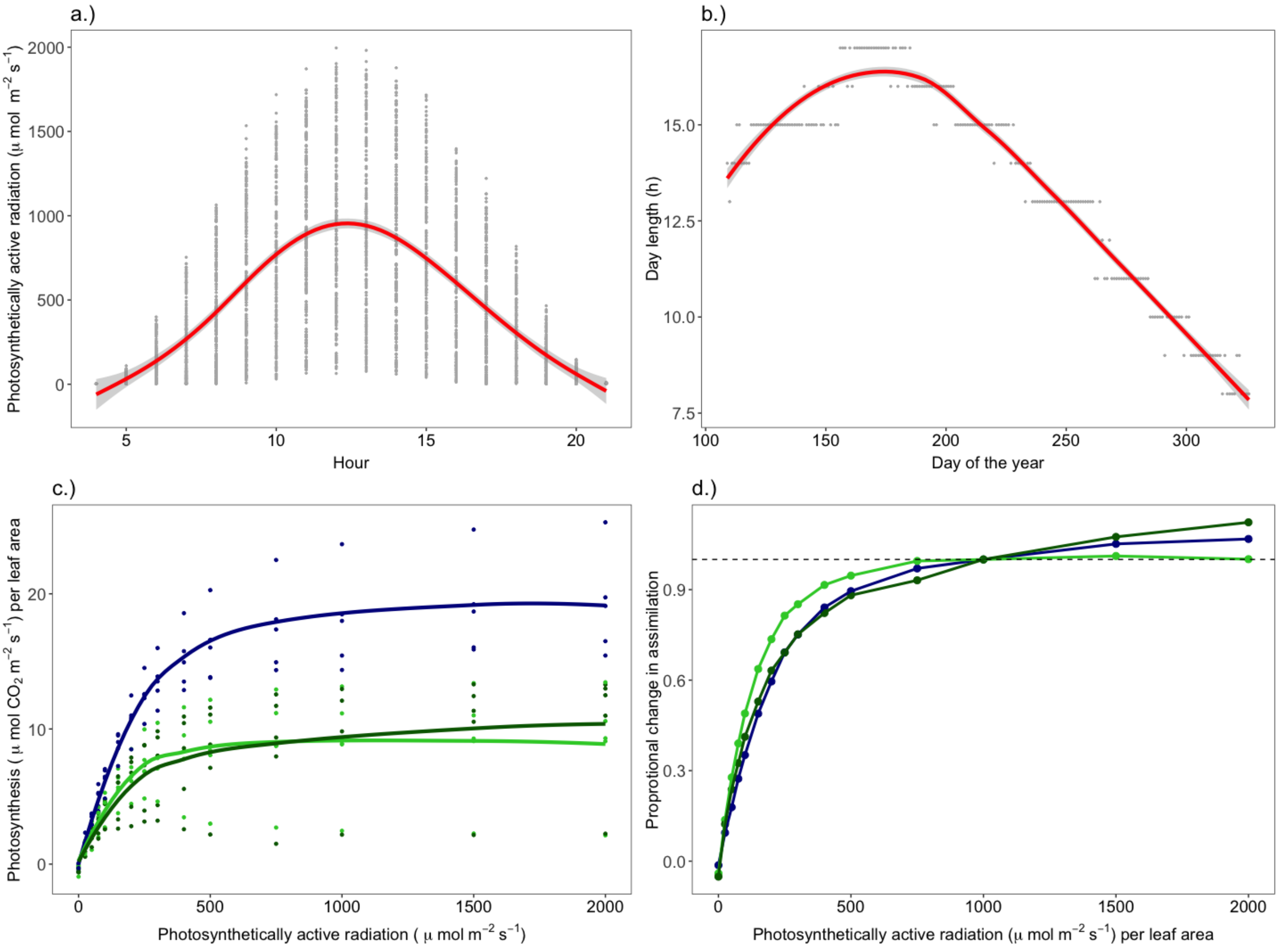


**Figure S2.** The underlying data and assumptions for the model on canopy photosynthesis. a) The diurnal change in solar radiation for each day during the growing season (Rennie et al. 2017). The hourly measurements are shown as grey dots, and the red line represents a smoothed average. b) The length of the day (h) over the growing season. c) The photosynthetic response of oak leaves to light (Visakorpi et al. 2018) for each of the three leaf types separately. d) The proportional change in photosynthesis when compared to the photosynthetic rate at light intensity 1000 µmol m$^{-2}$ s$^{-1}$ of photosynthetically active radiation. The dashed line shows the rate at light intensity 1000 µmol m$^{-2}$ s$^{-1}$ PAR, which was se as the level for 100% photosynthesis. In panels c) and d), the dark blue line shows the intact leaf, the light green the leaf damaged by herbivory and dark green the systemically affected leaf (i.e. an intact leaf growing on a same shoot with damaged leaves).

<u>Night-time canopy respiration</u>. To estimate night-time canopy respiration, we measured leaf respiration on the five oaks during two nights in July 2015 (8$^{th}$ - 9$^{th}$ July and 22$^{nd}$ - 23$^{rd}$ July). Measurements were taken after sunset and before sunrise (10pm - 2am) with an infra-red gas

analyser (CIRAS-2, PPsystems, Hitchin, UK). Temperature was kept as ambient, flowrate at 200 ml $min^{-1}$, and $CO_2$ concentration at 400 ppm. From each tree, we measured one completely intact leaf, one leaf damaged by herbivores and one intact leaf growing next to the damaged leaves, using the same experimental setup as for the photosynthetic measurements (Visakorpi et al. 2018). Since nightly respiration rates are usually very low, and therefore affected by possible leaks in the measuring chamber of the gas analyser, we measured a piece of paper at the beginning and at the end of each measuring night. If the rate measured from the paper was different from zero, this value was subtracted from the leaf respiration rates. All positive respiration values (apparent net $CO_2$ uptake) were discarded as measurement errors. Because the respiration rates did not show any trends throughout the night (Figure S3a), and because previous studies (Stokes 2002) have also shown that the respiration rate does not change throughout the night, we used mean respiration values per leaf type per tree for upscaling canopy respiration.

To estimate respiration under normal level of herbivory, the rates per unit leaf area were scaled with the proportion of the different leaf types in the canopy using the following equation (Visakorpi et al. 2018):

$$CR_{herb} = \sum_{t=1}^{3} R_t * (1 - D_t) * l_t,$$

(Eq. 4)

where $CR_{herb}$ is canopy respiration under herbivory, t denotes the three different leaf types (1 = completely intact, 2 = systemically affected, 3 = damaged), $D_t$ is the proportion of leaf area loss per leaf type, $R_t$ is the mean night-time respiration rate of that leaf type as µmol $CO_2$ $m^{-2}$ $d^{-1}$ $s^{-1}$, and $l_t$ is the proportion of leaf type $t$ in the canopy. The output is instantaneous canopy respiration as µmol $CO_2$ $m^{-2}$ $s^{-1}$ of leaf area at the top of the canopy. For an estimate of respiration for a completely intact canopy, we used the average value measured for the completely intact leaves.

Because night respiration often follows the total photosynthetic rate of the day before (Whitehead et al. 2004), we scaled the night-time respiration rates with a previously identified relationship between daytime canopy photosynthesis and respiration the following night (Whitehead et al. 2004):

$$R_t = 0.074\,A_t + 0.03,$$

(Eq. 5)

where $R_t$ (mol CO2 $m^{-2}$ $d^{-1}$) is total night time canopy respiration and $A_t$ (mol $CO_2$ $m^{-2}$ $d^{-1}$) is total daytime canopy photosynthesis. We estimated $R_t$ for each night based on the canopy photosynthesis of the previous day. These values were turned into proportional change in respiration from the reference night when the respiration measurements were taken ($8^{th}$ - $9^{th}$ July and $22^{nd}$ - $23^{rd}$ July). The canopy respiration rates estimated above (*CR*) were then scaled with this estimated proportional change over the season. We estimated and errors of 15% for this relationship, based on the original data (Whitehead et al. 2004).

To take into account the change in respiration rate though the different canopy layers, we used formula by (Griffin et al. 2001), who studied the change in nightly respiration rates through different canopy layers on red beech (*Nothofagus fusca,* Hook. F.). We calculated respiration rate per unit leaf area for each one meter layer in the canopy, using the average canopy height for oaks measured at our study site (7.8 m, see above "Tree height and crown area") and the following equation:

$$CR_L = -0.0028 * height + CR,$$

(Eq. 6)

where $CR_L$ is canopy respiration per canopy layer $L$, *height* is the height of the layer in meters from the top of the canopy and $CR$ is the canopy respiration measured at the top of the canopy (µmol $CO_2$ $m^{-2}$ $d^{-1}$ $s^{-1}$). We divided each value with the average canopy height and summed these estimates together to yield an estimate of the average respiration rate per unit leaf area of the whole canopy.

Since variation in respiration rate is mainly controlled by temperature (Ryan 1991, Tjoelker et al. 2001), we scaled the canopy respiration rates with the ambient air temperatures measured in Wytham Woods throughout the season of 2015 (Figure S3b) with the following equation (Bolstad et al. 2003):

$$R_T = R_{Tref} * Q_{10}^{\frac{(T-T_{ref})}{10}},$$

(Eq. 7)

where $R_T$ is respiration rate at a given temperature $T$ (ºC), $R_{Tref}$ is the respiration rate at reference temperature $T_{ref}$, and $Q_{10}$ is a coefficient describing ratio of respiration measured over a 10-degree span. As a reference temperature for the intact canopy, we used the average measuring temperature of the completely intact leaf on each tree. As a reference temperature for the canopy under normal levels of herbivory, we weighted the measuring temperatures of the three leaf types with their average proportion in the canopy and used the summed temperature as the reference temperature. Night-time hours were defined as those when light level was zero.

Since the temperature response of respiration differs between different canopy layers, we applied temperature correction for three canopy layers separately (top of the canopy 0-2 m, middle of the canopy 3-5 m and bottom of the canopy 6-8 m). We used average $Q_{10}$ values per canopy layer measured for oaks *Q. rubra* and *Q. prinus* (bottom layer $Q_{10} = 2.15 \pm 0.12$, middle layer $Q_{10} = 1.89 \pm 0.1$, top layer $Q_{10} = 1.87 \pm 0.19$; (Turnbull et al. 2003, Figure S3c), divided each estimate by three and summed these proportional estimates. This gave us respiration estimates for each hour for each night during the growing season for the whole canopy. We then summed the hourly estimates for each night (as area under curve) to achieve the respiration per night as µmol $CO_2$ $m^{-2}$ $d^{-1}$ of leaf area. Finally, this rate was multiplied with the LAI to achieve µmol $CO_2$ $m^{-2}$ $d^{-1}$ of ground area. Since scaling the respiration rates with the canopy photosynthesis of the previous day already includes the effect of reduced LAI in the spring and autumn, we did not apply any further corrections for the seasonal changes in LAI. The values for each night were summed to achieve the yearly $CO_2$ loss to night-time respiration per $m^2$ of ground area.

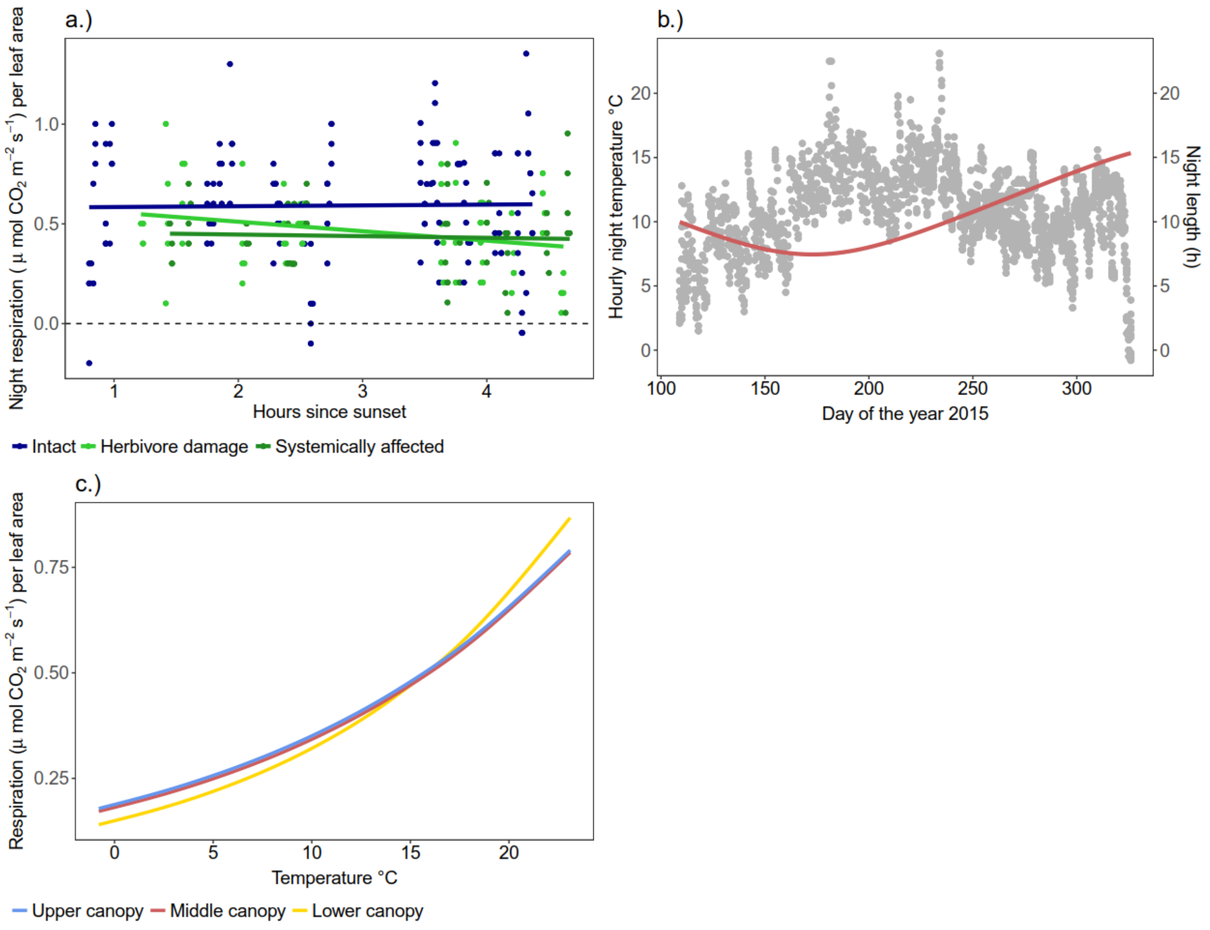


**Figure S3.** The underlying data and assumptions for the model on nightly canopy respiration. a) The measurements of night-time leaf respiration rates during summer 2015 at the upper canopy leaves in Wytham Woods. Negative values (i.e. carbon uptake) were deleted from the data set.  b) Hourly night temperatures during the summer 2015 in Wytham Woods (grey dots), and the length of the night in hours (red line). c) The response of leaf respiration rate to temperature for the three canopy layers (Turnbull et al. 2003). Note that the values have not been corrected to take into account the dependence of night-time respiration on the daytime canopy photosynthesis to make the relationship between temperature and respiration clearer.

Stem and root respiration. To estimate stem respiration, we used the average stem respiration rate measured in Wytham Woods during the summer 2017 from 25 mature oak trees at 1.3m with closed chamber method (Walker 2017) for the method, see (Marthews et al. 2014). One outlier measurement with a high stem respiration rate (26.6 g $CO_2$ $m^{-2}$ $d^{-1}$) was removed from the data set. Stem respiration was corrected for hourly temperature of the whole year, both day and night with Equation 7 (see "Leaf respiration" above). We assumed a $Q_{10}$ value of 2 (describing change in rate by 2 units for every 10 degrees), which is commonly used to describe the temperature response of stem respiration of temperate deciduous trees (e.g. $Q_{10}$ = 1.7 for beech, Damesin et al. 2002, $Q_{10}$ = 2.3 for *Q. rubra*; Bolstad et al. 2004). Since the differences between stem and air temperatures were small, we assumed stem temperature to equal air temperature. From the hourly data, we estimated carbon loss per day per unit stem area. The temperature corrected hourly stem respiration rates were then summed to achieve the amount of $CO_2$ respired by each stem in 24h over the year. Since stem respiration shows a positive relationship with leaf photosynthesis similar to leaf respiration (Wertin and Teskey 2008), we assumed that increased photosynthesis in the absence of herbivores would also increase stem respiration rates. To model this, we calculated the ratio between leaf respiration between intact and normal canopies and used this to scale respiration measures for the intact canopy (Table S2).

To estimate root respiration, we calculated the ratio between stem and root respiration based on earlier measurements at the plot (Fenn et al. 2015). This ratio (root respiration 13.0 ± 4.0 % of stem respiration) was used to scale the estimated stem respiration values into root respiration values.

**Table S2.** Allometric relationships on oak at the study site, used for estimating total oak net primary productivity (NPP): NPP of leaves and wood, above-ground NPP (ANPP; the sum of leaf and wood NPP), the proportional allocation of ANPP into wood production, ratio of below-ground and above-ground NPP and ratios of leaf to wood NPP, intact to normal canopy gas exchange and canopy respiration and root to stem respiration.

| Measure | Estimate | SEM | Reference |
|---|---|---|---|
| Oak leaf NPP, Mg C ha$^{-1}$ yr$^{-1}$ | 0.25 | 0.14 | Fenn 2010 |
| Oak wood NPP, Mg C ha$^{-1}$ yr$^{-1}$ | 0.3 | 0.07 | Fenn 2010 |
| Oak ANPP, Mg C ha$^{-1}$ yr$^{-1}$ | 0.55 | 0.1 | Fenn 2010 |
| Allocation of NPP into wood | 55% | 20% | Fenn 2010 |
| Ratio of leaf NPP to wood NPP | 0.85 | 0.31 | Fenn 2010 |
| NPP below/NPP above | 0.59 | 0.18 | Fenn 2010 |
| Ratio of intact to normal canopy gas exchange | 2.19 | 0.51 | This study |
| Ratio of intact to normal respiration | 1.42 | 0.23 | This study |
| Ratio of root respiration to stem respiration | 0.13 | 0.04 | (Fenn et al. 2015) |

Combining stem, root and canopy estimates. To combine estimates of tree-level stem and root respiration with the canopy gas exchange and to scale these estimates for a hypothetical hectare of the forest comprised of oak trees only (although in reality the site is a mixed species forest), we assumed LAI of 6.5 m$^2$/m$^2$ and stem area index ("SAI") of 1.5 m$^2$/m$^2$. The LAI is based on previous litter fall data at the site (Fenn 2010, Nils Rutjes 2016, unpublished data), and SAI is based on estimates from other forests with similar age and structure (Fenn et al. 2015). We assumed a 50% error for the SAI estimate to reflect the uncertainty in this estimate (Fenn et al. 2015). NPP was estimated by subtracting the sum of leaf, stem and root respiration from photosynthesis. To estimate oak NPP per hectare using the actual density of the oak trees in the plot (20% of the total plot basal area, Table 1 in the main text), we multiplied the values per m$^2$ of basal area with the oak basal area of the plot. Finally, the amount of $CO_2$ photosynthesised or respired, measured as micromoles, were converted to grams of carbon by multiplying the rate as moles with the molar mass of $CO_2$ (44.01) and with the ratio of carbon molar mass to $CO_2$ molar mass (0.27).

*2.) NPP through tree growth census*

To compare the estimate of oak NPP derived through canopy upscaling with NPP estimates obtained through other methods, we used data on tree dbh measures on all oak trees (355 stems) within the 18-ha plot (see "Study site" in the main text) collected in 2010 and 2016 (Y. Malhi, unpublished analysis). We then used the following equation (Fenn et al. 2015) to estimate the aboveground oak woody biomass:

$$ABW = e^{-5.284602+(2.4682\ln c)},$$

(Eq. 8)

where *ABW* is the aboveground woody biomass of oak (as kg) and *c* is the circumference of the tree (in cm) at breast height (Bunce 1968). We estimated *ABW* for all oak trees for both censuses, converted biomass into carbon by multiplying with the carbon content of oak wood (47% ± 1%; (Butt et al. 2009) and used the change in above-ground woody carbon stock between 2010 and 2016 as an estimate of woody NPP. The change in ABW was divided by the length of the census interval (6 years) to estimate yearly growth in ABW. We then used oak-specific allometric relationships measured previously at the plot to estimate oak canopy NPP and belowground NPP (Fenn 2010); Table S2). To estimate the effect of herbivory on oak NPP, we estimated NPP in the absence of herbivory assuming that herbivory reduced total NPP to the same extent as canopy gas exchange in the canopy upscaling calculations described above (54%, Table S2). To estimate oak NPP per ha of the actual study site, we summed NPP estimates of each tree in the study area and divided this with the size of the surveyed area (18 ha). To estimate NPP per hectare in hypothetical forest comprised only of oak trees, we divided the woody NPP per ha of the real plot by the contribution of oak to the total plot basal area (Table 1 in the main text) and the leaf NPP by the contribution of oak to the total plot LAI (Table 1 in the main text). Trees with large negative growth rates (larger than 1.0 cm reduction in dbh per year, n = 6) and trees that had died during the census interval (n = 12) were removed from the data set. There was no oak recruitment during the census interval. The final errors represent sampling error in tree circumference change (n = 355), the error in the oak wood C content, and the uncertainties in the relationship between aboveground to belowground NPP, wood NPP to leaf NPP and intact to normal canopy gas exchange (Table S2).

*3.) NPP through earlier biometric measurements*

To obtain a third estimate on the effect of herbivory on oak NPP, we used an earlier estimate for oak NPP at the plot based on biometric measures (Fenn 2010, Fenn et al. 2015). These measures include aboveground oak NPP for an area of one hectare within the 18-ha plot, oak-specific ratio of wood NPP to leaf NPP and oak-specific ratio of aboveground biomass to belowground biomass (Table S2). We combined these relationships to obtain an estimate of oak NPP as Mg C $yr^{-1}$ per ha for the specific one hectare subplot in which the original study by Fenn and others (2015) took place. To extrapolate the results to a wider 18-ha study site, the estimate was divided by the oak basal area of the one hectare study plot, and multiplied by the mean oak basal area of the whole site (6.7 $m^2$ $ha^{-1}$, Table 1 in the main text). Similarly, to estimate oak NPP per hectare of a hypothetical forest consisting only of oak trees, we divided the estimate by the oak basal area of the studied one hectare plot, and multiplied the estimate by the total basal area of the plot (33 $m^2$ $ha^{-1}$, Table 1 in the main text). To estimate the effect of herbivory on oak NPP, we again assumed that the total NPP was reduced to the same extent as canopy gas exchange in the canopy upscaling calculations above. The error here represents the propagated uncertainties with the original oak NPP estimate, the ratio of aboveground to belowground NPP, wood NPP to leaf NPP and intact to normal canopy gas exchange (Table S2).

**Error propagation**

The uncertainty in the daytime net canopy photosynthesis values was estimated by combining the sampling error in the original photosynthesis measurements and in the proportion of the different leaf types (8% for an intact canopy, 13% for a normal canopy), the mean error in the seasonal change in photosynthesis, estimated from the original data (Morecroft et al. 2003, 23%), the mean error in LAI change, estimated from the error in expanding leaves in the spring (Stokes 2000, 0.5%) and the mean error in photosynthetic response to light (6%), estimated as a GAM based on the light-response curves (see above, "Diurnal pattern in photosynthesis").

The uncertainty in the night-time canopy respiration values was estimated by combining the sampling error in the original respiration measurements and in the proportion of the different leaf types (5% for an intact canopy, 7% for a normal canopy) and the mean error in the relationship between daily photosynthesis and nightly respiration (Whitehead et al. 2004, 15%).

The error for stem respiration was estimated by combining the sampling error in the original stem respiration measurements (Walker 2017, 40%) and the estimated error in SAI (Fenn et al. 2015, 50%). The error for root respiration was estimated by combining error in stem respiration measurement and the uncertainty in scaling from stem to root respiration (31%). For the estimate of the woody respiration of an intact tree, the error also includes the uncertainty in scaling from normal canopy to an intact canopy (Table S2, 23%).

For estimating the error for the total NPP, we first estimated error for canopy respiration without taking into account for errors associated with the proportion of the different leaf types in the canopy, and for root respiration without taking into account the uncertainty in SAI or the raw stem respiration measurements. This was done to avoid accounting for these uncertainties twice in the final NPP estimate.

To estimate the error for the differences in NPPs in the two herbivory scenarios (*Intact canopy* and *Normal herbivory*, i.e. the herbivory effect on NPP), we combined measurement errors of the instantaneous canopy photosynthesis and respiration between intact canopies and canopies experiencing normal level of herbivory. This was done to avoid including uncertainty associated with the different scaling functions (e.g. photosynthesis to light, or seasonality in LAI) twice (i.e. as associated with the individual estimates for NPP in the two scenarios).

Errors were combined using standard error propagation rules: when summing values that each had an associated error, the errors were squared and summed, and from the resulting sum we took a square root. When values with errors were multiplied or divided, the proportional errors were summed, squared, and from the resulting sum we took a square root.

All analyses were carried out using R version 3.5.0, (R Core Team 2018) and packages geosphere (for daylength estimation; Hijmans 2017), MESS (for calculating area under curve, Ekstrøm 2018), and mgcv (for GAM models; Wood 2011).

**References for Appendix 1**

Bolstad, P. V., K. J. Davis, J. Martin, B. D. Cook, and W. Wang. 2004. Component and whole-system respiration fluxes in northern deciduous forests. Tree Physiology 24:493–504.

Bolstad, P. V., P. Reich, and T. Lee. 2003. Rapid temperature acclimation of leaf respiration rates in Quercus alba and Quercus rubra. Tree Physiology 23:969–976.

Bunce, R. 1968. Biomass and production of trees in a mixed deciduous woodland: I. Girth and height as parameters for the estimation of tree dry weight. Journal of Ecology 5:759–757.

Butt, N., G. Campbell, Y. Malhi, M. Morecroft, K. Fenn, and M. Thomas. 2009. Initial results from establishment of a long-term broadleaf monitoring plot at Wytham Woods, Oxford, UK. University of Oxford, Oxford.

Clark, D. B., L. M. Mercado, S. Sitch, C. D. Jones, N. Gedney, M. J. Best, M. Pryor, G. G. Rooney, R. L. H. Essery, E. Blyth, O. Boucher, R. J. Harding, C. Huntingford, and P. M. Cox. 2011.

The Joint UK Land Environment Simulator (JULES), model description – Part 2: Carbon fluxes and vegetation dynamics. Geoscientific Model Development 4:701–722.

Damesin, C., E. Ceschia, N. Le Goff, J.-M. Ottorini, and E. Dufrêne. 2002. Stem and branch respiration of beech: from tree measurements to estimations at the stand level. New Phytologist 153:159–172.

Ekstrøm, C. T. 2018. MESS: Miscellaneous esoteric statistical scripts.

Fenn, K. M. 2010. Carbon cycling in British deciduous woodland: processes, budgets, climate & phenology. DPhil, University of Oxford.

Fenn, K., Y. Malhi, M. Morecroft, C. Lloyd, and M. Thomas. 2015. The carbon cycle of a maritime ancient temperate broadleaved woodland at seasonal and annual scales. Ecosystems 18:1–15.

Griffin, K. L., D. T. Tissue, M. H. Turnbull, W. Schuster, and D. Whitehead. 2001. Leaf dark respiration as a function of canopy position in Nothofagus fusca trees grown at ambient and elevated CO2 partial pressures for 5 years. Functional Ecology 15:497–505.

Hijmans, R. J. 2017. geosphere: Spherical Trigonometry.

Kok, B. 1956. On the inhibition of photosynthesis by intense light. Biochimica et Biophysica Acta 21:234–244.

Marino, G., M. Aqil, and B. Shipley. 2010. The leaf economics spectrum and the prediction of photosynthetic light-response curves: Leaf economics spectrum and light-response. Functional Ecology 24:263–272.

Marthews, T., T. Riutta, I. Oliveras Menor, R. Urrutia, S. Moore, D. B. Metcalfe, Y. Malhi, O. L. Phillips, W. Huraca Huasco, M. Ruiz Jaén, C. Girardin, N. Butt, and R. Cain. 2014. Measuring Tropical Forest Carbon Allocation and Cycling: A RAINFOR-GEM Field Manual for Intensive Census Plots (v3.0). Manual.

Mercado, L. M., C. Huntingford, J. H. C. Gash, P. M. Cox, and V. Jogireddy. 2007. Improving the representation of radiation interception and photosynthesis for climate model applications. Tellus B: Chemical and Physical Meteorology 59:553–565.

Monsi, M., and T. Saeki. 1953. Ueber den Lichtfaktor in den Planzengesellschaften und seine Bedeutung fuer die Stoffproduktion. Journal of Japanese Botany:22–52.

Morecroft, M. D., and J. M. Roberts. 1999. Photosynthesis and stomatal conductance of mature canopy Oak (Quercus robur) and Sycamore (Acer pseudoplatanus) trees throughout the growing season. Functional Ecology 13:332–342.

Morecroft, M. D., V. J. Stokes, and J. I. L. Morison. 2003. Seasonal changes in the photosynthetic capacity of canopy oak (Quercus robur) leaves: the impact of slow development on annual carbon uptake. International Journal of Biometeorology 47:221–226.

R Core Team. 2018. R: A language and environment for statistical computing. R Foundation for Statistical Computing, Vienna, Austria.

Rennie, S., J. Adamson, R. Anderson, C. Andrews, J. Bater, N. Bayfield, K. Beaton, D. Beaumont, S. Benham, V. Bowmaker, C. Britt, R. Brooker, D. Brooks, J. Brunt, G. Common, R. Cooper, S. Corbett, N. Critchley, P. Dennis, J. Dick, B. Dodd, N. Dodd, N. Donovan, J. Easter, E. Eaton, M. Flexen, A. Gardiner, D. Hamilton, P. Hargreaves, M. Hatton-Ellis, M. Howe, J. Kahl, M. Lane, S. Langan, D. Lloyd, B. McCarney, Y. McElarney, C. McKenna, S. McMillan, F. Milne, L. Milne, M. Morecroft, M. Murphy, A. Nelson, H. Nicholson, D. Pallett, D. Parry, I. Pearce, G. Pozsgai, R. Rose, S. Schafer, T. Scott, L. Sherrin, C. Shortall, R. Smith, P. Smith, R. Tait, C. Taylor, M. Taylor, M. Thurlow, A. Turner, K. Tyson, H. Watson, M. Whittaker, M. Wilkinson, and C. Wood. 2017. UK Environmental Change Network (ECN) meteorology data: 1991-2015. NERC Environmental Information Data Centre.

Ryan, M. G. 1991. Effects of climate change on plant respiration. Ecological Applications 1:157–167.

Sager, J. C., and C. McFarlane. 1997. Radiation. Page *in* R. W. Langhans and T. W. Tibbitts, editors. Plant growth chamber handbook. Iowa, U.S.A.

Stokes, V. 2000. Effects of microenvironment and leaf developmental characteristics on annual carbon gain and water use in two deciduous tree species. DPhil, University of Oxford.

Tjoelker, M. G., J. Oleksyn, and P. B. Reich. 2001. Modelling respiration of vegetation: evidence for a general temperature-dependent Q10. Global Change Biology 7:223–230.

Turnbull, M. H., D. Whitehead, D. T. Tissue, W. S. F. Schuster, K. J. Brown, and K. L. Griffin. 2003. Scaling foliar respiration in two contrasting forest canopies. Functional Ecology 17:101–114.

Visakorpi, K., S. Gripenberg, Y. Malhi, C. Bolas, I. Oliveras, N. Harris, S. Rifai, and T. Riutta. 2018. Small-scale indirect plant responses to insect herbivory could have major impacts on canopy photosynthesis and isoprene emission. New Phytologist 220.

Visakorpi, K., T. Riutta, Y. Malhi, J.-P. Salminen, N. Salinas, and S. Gripenberg. 2020. Changes in oak (Quercus robur) photosynthesis after winter moth (Operophtera brumata) herbivory are not explained by changes in chemical or structural leaf traits. PLOS ONE 15:e0228157.

Walker, K. 2017. Variation and patterns of $CO_2$ efflux and sapwood content of three deciduous broadleaved trees. B.S. thesis, University of Oxford, Oxford, UK.

Wertin, T. M., and R. O. Teskey. 2008. Close coupling of whole-plant respiration to net photosynthesis and carbohydrates. Tree Physiology 28:1831–1840.

Whitehead, D., K. L. Griffin, M. H. Turnbull, D. T. Tissue, V. C. Engel, K. J. Brown, W. S. F. Schuster, and A. S. Walcroft. 2004. Response of total night-time respiration to differences in total daily photosynthesis for leaves in a *Quercus rubra* L. canopy: implications for modelling canopy $CO_2$ exchange. Global Change Biology 10:925–938.

Wood, S. N. 2011. Fast stable restricted maximum likelihood and marginal likelihood estimation of semiparametric generalized linear models: Estimation of Semiparametric Generalized Linear Models. Journal of the Royal Statistical Society: Series B (Statistical Methodology) 73:3–36.
