## Appendix 2 for "Does insect herbivory suppress ecosystem productivity? Evidence from a temperate woodland"

### Appendix 2: Additional figures and tables

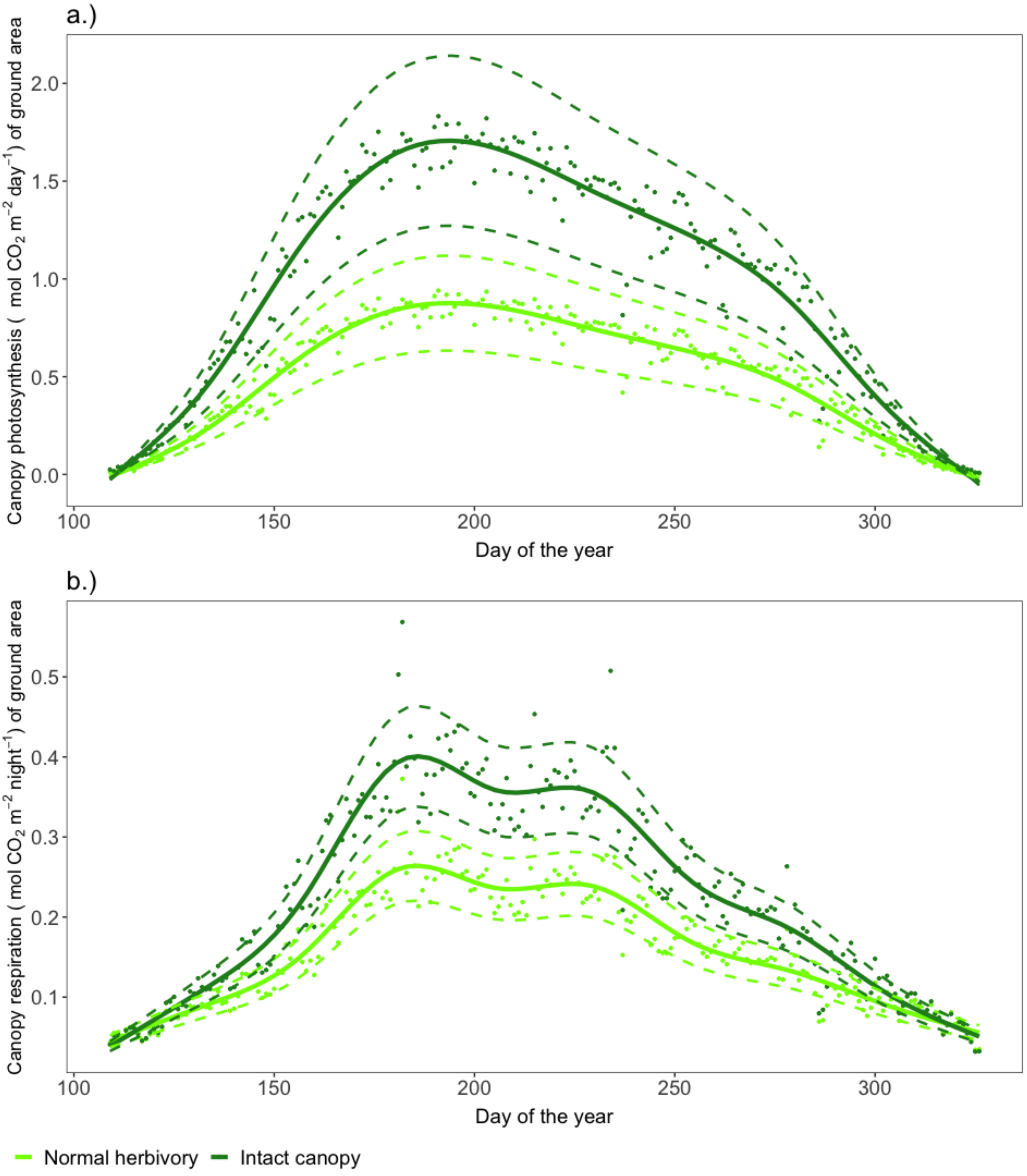


**Figure S5**. a) Daily canopy net photosynthesis and b) night-time canopy respiration per $m^2$ of ground area based on data from five oak trees in Wytham Woods during the growing season 2015 ($19^{th}$ April to $22^{nd}$ November). The model for canopy photosynthesis takes into account the effect of reduced light through the canopy layers and daytime respiration rate per unit leaf area (Visakorpi et al. 2018), seasonality in photosynthetic rate per unit leaf area (Morecroft et al. 2003), seasonality in LAI, change in light intensity during each day and daylength over the season, and photosynthetic response to light (Visakorpi et al. 2018). The “intact canopy” scenario is based on measurements on intact leaves surrounded by only intact leaves, whereas the estimates under “normal herbivory” are on photosynthesis and respiration rates of three different leaf types (intact, damaged, systemically affected), weighted with their observed abundance. The canopy respiration model takes into account different respiration rates at different canopy layers, the effect of temperature on respiration scaled for hourly temperature data, the differences in temperature responses at different part of the canopy, the effect of daytime photosynthesis on night-time respiration and seasonality of LAI. In all panels, each data point is a modelled estimate for the average rate of the five trees for one day. The solid line shows a general additive model across the five trees, and the dashed coloured lines represent the propagated measuring uncertainty.

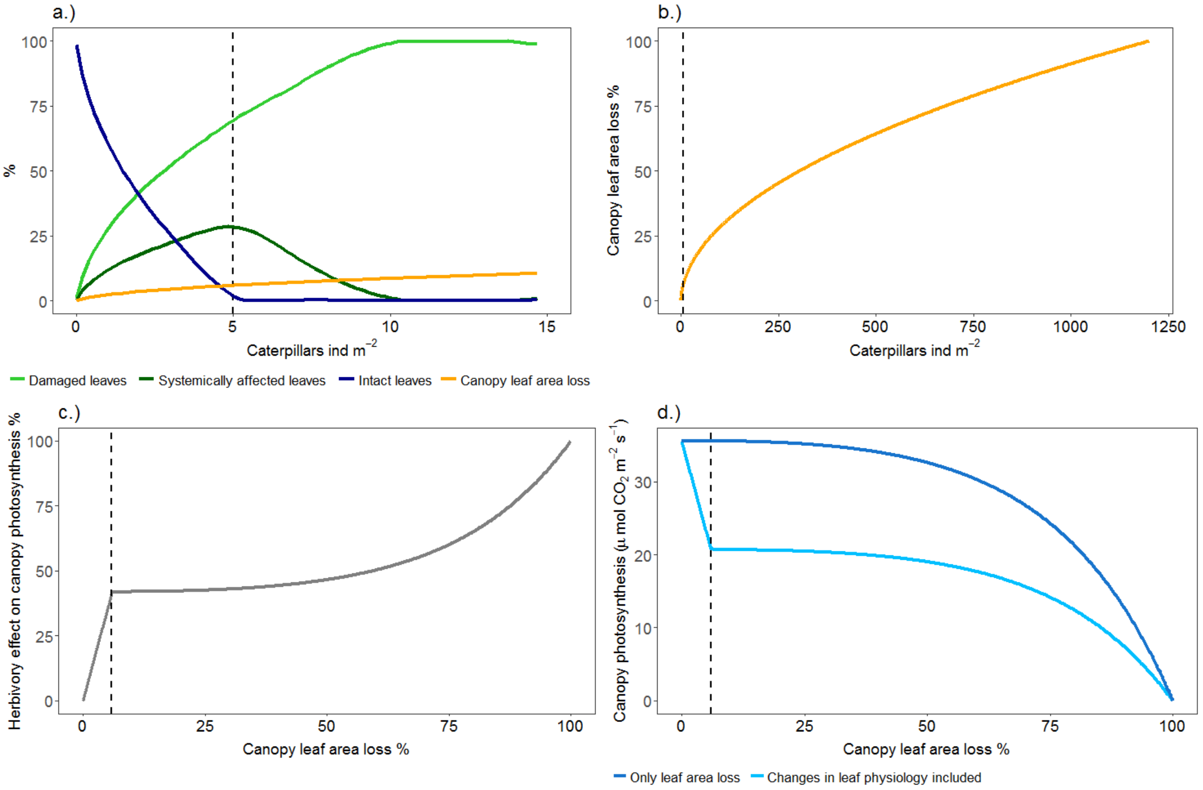


**Figure S6.** Simulated proportions of the three different leaf types (intact, systemically affected and damaged) and leaf area loss in the canopy with increasing winter moth caterpillar density and the effect of increasing herbivory on canopy photosynthesis. The dashed line denotes the conditions during our sampling year, 2015 (5 caterpillar individuals m$^{-2}$ and 5.9% leaf area loss). a) Caterpillar density up to 15 individuals m$^{-2}$ and the proportion of the three different leaf types (intact, systemically affected, damaged) and leaf area loss. b) The relationship between caterpillar density and leaf area loss until complete defoliation. c) The effect of increasing herbivory on canopy photosynthesis (% of reduction from the full potential rate), and d) canopy photosynthesis with increasing leaf area loss in two scenarios: when only the effect of leaf area loss is taken into account, and when the herbivore-induced changes in leaf physiology are included in the canopy photosynthesis estimate.

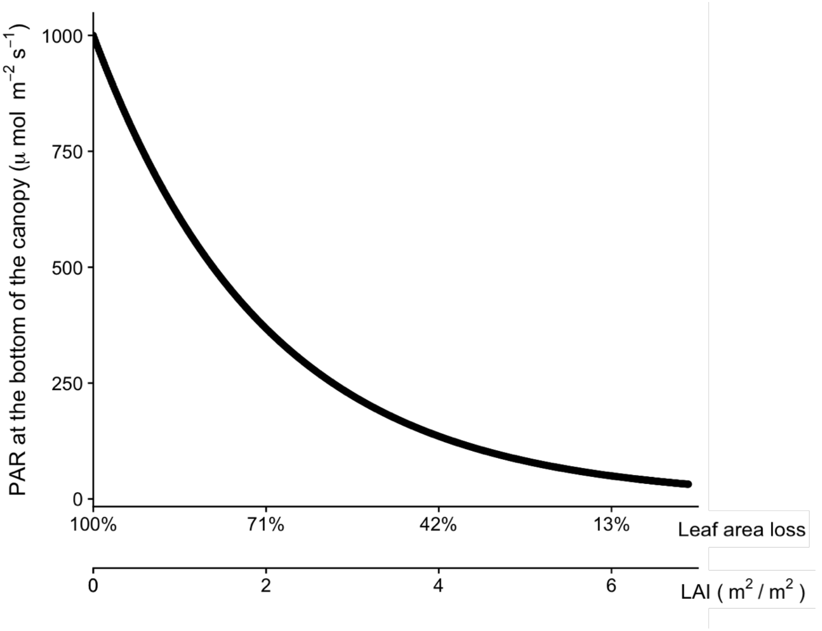


Figure S7. The relationship between leaf area index (LAI), leaf area loss and the amount of light at the bottom of the canopy when the light intensity is 1000 µmol m$^{-2}$ s$^{-1}$ PAR at the top.

**Table S3.** Leaf area loss to herbivory, mean leaf mass per area (LMA) and leaf carbon content in the five studied oak trees. The errors are ± 1 SEM representing the error over several leaf samples per tree. N is sample size (i.e. number of sampled leaves) per tree.

| Oak ID | % of herbivory | n | Mean LMA (gm$^{-2}$) | n | Mean C (%) of leaf | n |
|---|---|---|---|---|---|---|
| 1 | 4.58 ± 0.8 | 164 | 58.1 ± 4.5 | 11 | 44.6 ± 1.2 | 4 |
| 2 | 7.02 ± 0.8 | 166 | 64. 8± 2.8 | 43 | 50.9 ± 1.4 | 3 |
| 3 | 7.54 ± 0.9 | 199 | 73.0 ± 1.5 | 41 | 50.6 ± 1.0 | 4 |
| 4 | 4.84 ± 0.8 | 191 | 63.6 ± 1.4 | 100 | 46.3 ± 1.0 | 6 |
| 5 | 5.34 ± 0.9 | 126 | 60.1 ± 1.5 | 40 | 50.2 | 1 |

#### References for Appendix 2

Morecroft, M. D., V. J. Stokes, and J. I. L. Morison. 2003. Seasonal changes in the photosynthetic capacity of canopy oak (Quercus robur) leaves: the impact of slow development on annual carbon uptake. International Journal of Biometeorology 47:221–226.

Visakorpi, K., S. Gripenberg, Y. Malhi, C. Bolas, I. Oliveras, N. Harris, S. Rifai, and T. Riutta. 2018. Small-scale indirect plant responses to insect herbivory could have major impacts on canopy photosynthesis and isoprene emission. New Phytologist 220.
