## Appendix 3 for "Does insect herbivory suppress ecosystem productivity? Evidence from a temperate woodland"

### Appendix 3. The relationship between the effect of herbivory on canopy assimilation and the intensity of herbivory under different scenarios

We explored how sensitive our model-inferred relationships between the effect of herbivory on canopy assimilation (as the difference between an intact canopy and a canopy with herbivory) and the intensity of herbivory (as % of leaf area loss) were to our model assumptions by simulating the relationship under four alternative scenarios. First, we assumed that the  difference in photosynthetic rate between intact and damaged leaves would be smaller than what our field measurements (Visakorpi et al. 2018) suggest. Second, we tested a scenario in which the photosynthetic rate of intact leaves depends on the herbivory level of the whole tree. Third, we simulated a different pattern at which herbivory spreads through the canopy. Finally, we tested how an additional leaf flush would affect the simulated relationship between herbivory and productivity.

To test the first scenario, we assumed that the photosynthetic rate of intact leaves would be a third smaller than observed in the previous study (Visakorpi et al. 2018). Based on this scenario, the role of indirect effects is smaller (ca. 25%) compared to when the measured photosynthetic rates are used for the simulations (ca. 50%). The shape of the relationship remains the same (Figure S8a).

To test the second scenario, we assumed that photosynthetic rate of completely intact leaves depends on their frequency in the canopy and is highest when this leaf type is rare (i.e. when the level of herbivory in the canopy is high) (Figure S8b). This describes a situation in which intact leaves are not able to keep a high photosynthetic rate when their proportion in the canopy is high, and/or in which the photosynthetic rate of intact leaves  increases to compensate the negative effects of herbivory on photosynthesis (e.g. Thomson et al. 2003, Retuerto et al. 2004). To simulate this, we built a linear model on the relationship between leaf area loss and photosynthesis of the intact leaf per unit leaf area for our five study trees (Figure S9a), and used this relationship to predict photosynthetic rates of intact leaves under different rates of herbivory (Figure S9b). We set a lower threshold, so that the photosynthetic rate of an intact leaf would always be at least 10% higher than the photosynthetic rate of a systemically affected leaf. As a result, the relationship between leaf area loss and the effect of herbivory on canopy assimilation is similar to that when the difference between photosynthetic rates of intact and damaged leaves is assumed to be smaller: the role of indirect effects is smaller (ca. 25%) than when intact leaves are assumed to photosynthesise at constant rate (ca. 50%).

To teste the third scenario, we assumed that herbivory would not spread in the canopy evenly to all shoots before spreading to all leaves (Figure S8c). Instead, we assumed that herbivory spreads first to all leaves within a shoot, and then to neighbouring shoots. Once all leaves have been damaged, the leaf area loss per leaf starts to increase (Figure S9c). Changing this assumption does not result in changes in the magnitude of the effect of herbivory on net photosynthesis.

To test the fourth assumption, we assumed a second leaf flush (“lammas shoots” on oak) to occur on August $1^{st}$ (called “lammas day” in England), the new leaf area to correspond to 50% of the leaf area lost, and all the new leaves to remain intact (and thus to photosynthesize at higher rate compared to the damaged and systemically affected leaves). We calculated canopy assimilation over the season with and without the lammas leaves, assuming different levels of herbivory. To simplify the calculations, we only included the effects of changing LAI on canopy photosynthesis, and not the changes in light intensity or daylength. Including the second leaf flush reduced the effect of herbivory on canopy assimilation at high herbivory intensities (Figure S8d): for example, the effect of herbivory does not exceed 75% even with 100% leaf area loss (because 50% of leaves come back as intact from August $1^{st}$ onward).

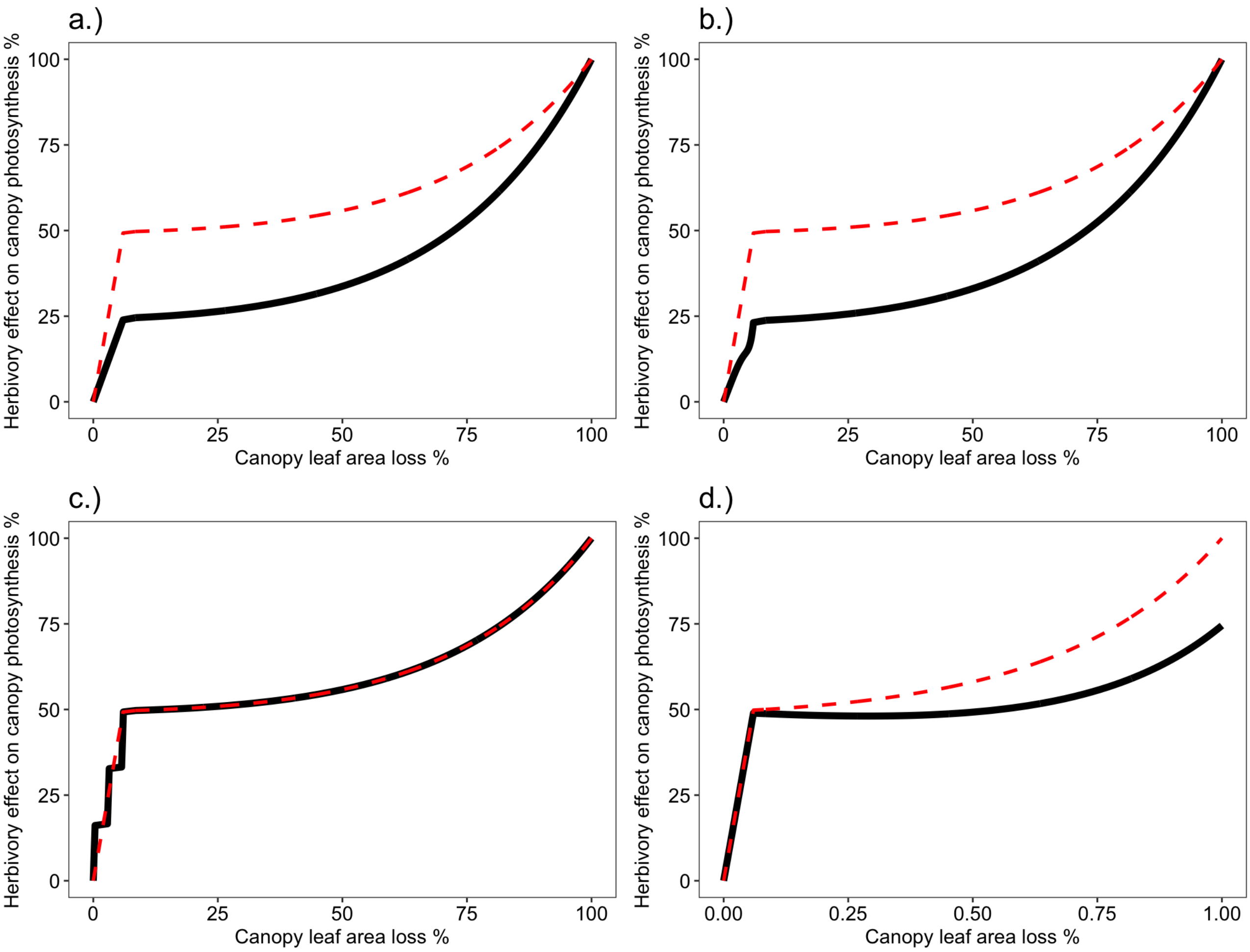


**Figure S8.** The effect of herbivory on canopy photosynthesis when a) the difference in photosynthetic rates between intact leaves and damaged leaves is assumed to be smaller than measured in our study (24% instead of 49%), b) when the photosynthetic rate of intact leaves is assumed to increase when their proportion in the canopy decreases, c) when herbivory is assumed to spread to each leaf within a shoot before spreading to neighbouring shoots (as opposed to spreading evenly to all shoots first) and d) when a second leaf flush is assumed to compensate 50% of the leaf area loss. The red dashed line represents the assumptions used in the study (corresponding to Figure 4 in the main text).

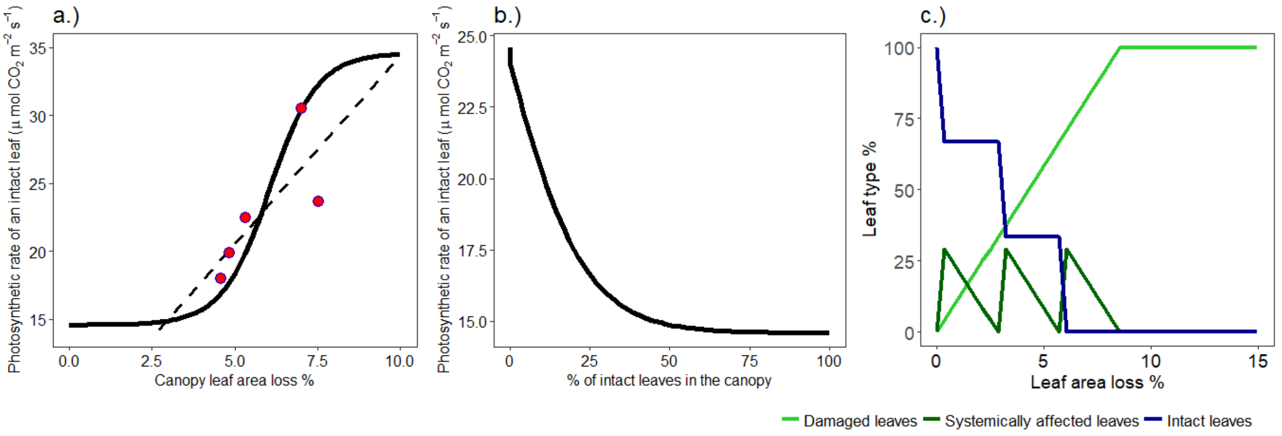


**Figure S9**. The model assumptions of the relationships described in Figure S8. Panel a) shows the photosynthetic rate per unit leaf area of intact leaf against percentage of leaf area loss in the canopy. Red dots show the measured rates of photosynthesis and herbivory in the five studied trees. The dashed black lines show a linear model drawn through these points, and black solid line shows the model used to predict the decrease in photosynthesis of intact leaves with decreasing leaf area loss (intact leaves perform the better the rarer they are). Panel b) shows the relationship between photosynthetic rate per unit leaf area of an intact leaf and the percentage of those leaves in the canopy, if the relationship in panel a (solid line) is assumed. Panel c) shows the proportions of the three different leaf types (damaged, systemically affected, intact) if herbivory is assumed to spread to all leaves within a shoot before spreading to new intact shoots.

#### References for Appendix 3

Retuerto, R., B. Fernandez-Lema, Rodriguez-Roiloa, and J. R. Obeso. 2004. Increased photosynthetic performance in holly trees infested by scale insects. Functional Ecology 18:664–669.

Thomson, V., S. Cunnigham, M. Ball, and A. Nicotra. 2003. Compensation for herbivory by Cucumis sativus through increased photosynthetic capacity and efficiency. Oecologia 134:167–175.

Visakorpi, K., S. Gripenberg, Y. Malhi, C. Bolas, I. Oliveras, N. Harris, S. Rifai, and T. Riutta. 2018. Small-scale indirect plant responses to insect herbivory could have major impacts on canopy photosynthesis and isoprene emission. New Phytologist 220.
